# Non-Poissonian bursts in the arrival of phenotypic variation can strongly affect the dynamics of adaptation

**DOI:** 10.1101/2023.11.02.565172

**Authors:** Nora S. Martin, Steffen Schaper, Chico Q. Camargo, Ard A. Louis

## Abstract

The introduction of novel phenotypic variation in a population through random mutations plays a crucial role in evolutionary dynamics. Here we show that, when the probability that a sequence has a particular phenotype in its 1-mutational neighbourhood is low, statistical fluctuations imply that in the weak-mutation or monomorphic regime, novel phenotypic variation is not introduced at a constant rate, but rather in non-Poissonian “bursts”. In other words, a novel phenotype appears multiple times in quick succession, or not at all for many generations. We use the RNA secondary-structure genotype-phenotype map to explore how increasing levels of heterogeneity in mutational neighbourhoods strengthen the bursts. Similar results are obtained for the HP model for protein tertiary structure and the Biomorphs model for morphological development. Burst can profoundly affect adaptive dynamics. Most notably, they imply that differences in arrival rates of novel variation can influence fixation rates more than fitness differences do.

## INTRODUCTION

Darwinian evolution accomplishes change over time through the joint processes of variation and selection. There is a longstanding tradition that focuses on the second step of the evolutionary process, using population genetics calculations that describe how genetic drift and natural selection affect the fixation dynamics in a population that initially starts with multiple alleles with different fitness, but where no new alleles appear [1]. It also possible to include the first step in the evolutionary process, the introduction of novel phenotypic variation, within a population genetics framework [2]. Since the fitness value of an allele is fundamentally caused by the interaction of the phenotype it represents with the environment, one can think of alleles with different fitness as representing different phenotypes and introduce new alleles at random into the population. A common underlying assumption in this class of models is that the introduction of new alleles can be characterised by an average rate (see e.g. refs [2, 3]). While these rates can differ, these models assume that individual introductions are uncorrelated, leading to Poisson statistics. We will call models that make such assumptions *average-rate models*.

A more sophisticated way to treat the introduction of novel phenotypic variation is to consider a genotype-phenotype (GP) map that explicitly models how random mutations lead to new phenotypes [5–7]. Examples of well-studied GP maps include RNA secondary structures [8–11], simplified models of protein structure, such as the HP model of tertiary structure [12] and the polyomino model of protein quaternary structure [13] and gene regulatory networks [14–16]. These molecular systems are all relatively tractable, which facilitates comprehensive studies. At present there are not many GP maps for larger-scale phenotypes that allow for such detailed investigations. One exception is the GP map underlying Richard Dawkins’ biomorphs [17, 18] a simple model of development, which was recently shown to share many features with molecular GP maps [19].

Interestingly, despite the diversity of the biological phenomena they represent, all these GP maps exhibit certain common patterns [6, 7]. For example, they take into account the fact that many mutations are neutral, as emphasised by Kimura [20], i.e. the mappings are many-to-one [8, 21]. This allows the definition of neutral sets comprising all genotypes mapping to the same phenotype *p*. The sizes of these neutral sets typically differ by multiple orders of magnitude [6–8]. Neutral sets are typically highly connected through mutations, allowing them to be traversed via point mutations by an evolving population undergoing genetic drift [7]. This connectedness is only possible because two genotypes that are mutational neighbours are (much) more likely to map to the same phenotype than two randomly chosen genotypes are [22], part of a broader set of so-called “neutral correlations” [22]. There is a small caveat, namely that the whole neutral set may not be connected in this way, due in part to biophysical constraints (for example in RNA, one needs a double mutation to change a CG bond to a GC bond [23, 24]), so that the neutral set can be broken down into *neutral components* (NCs) that are fully connected.

One important motivation for including the complexity of a GP map is to study the dynamics of neutral evolution [10, 23, 25, 26], in particular because neutral sets are known to have highly heterogeneous structure. For example, different genotypes in the NC can have different robustness (i.e. a different number of neutral mutations per genotype) [9]. Several studies have shown that such inhomogeneities in a neutral set can affect populations undergoing neutral evolution. For example, differences in robustness within the NC can lead to an overdispersion in the rate of (neutral) fixations [25, 27–29]: Because the supply of neutral mutations is higher at genotypes with higher robustness, the time to a neutral fixation depends on the robustness of the current genotype and is therefore not constant.

The effect of structure in the GP map on the neutral evolutionary dynamics prompts the question of whether the inhomogeneities present in the GP map also shape the introduction of new phenotypes. To illustrate this point, Fig 1A shows a NC taken from the *L* = 12 RNA secondary structure GP map for a phenotype *p*_*g*_ (grey) and the point mutation links it makes to two other phenotypes, *p*_*r*_ (red) and *p*_*b*_ (blue). Each sequence or genotype with a particular novel phenotype *p*_*i*_ in its 1-mutational neighbourhood is a *portal genotype* 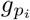 for *p*_*i*_ [30]. A population can only produce *p*_*i*_ as variation if it includes such portal genotypes 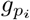 (except at very high mutation rates,when double mutations occur more frequently) and thus the distribution of portal genotypes over the NC is important for the dynamics of non-neutral variation. There are several reasons suggesting that this distribution of portals is not homogeneous.

**FIG. 1.**
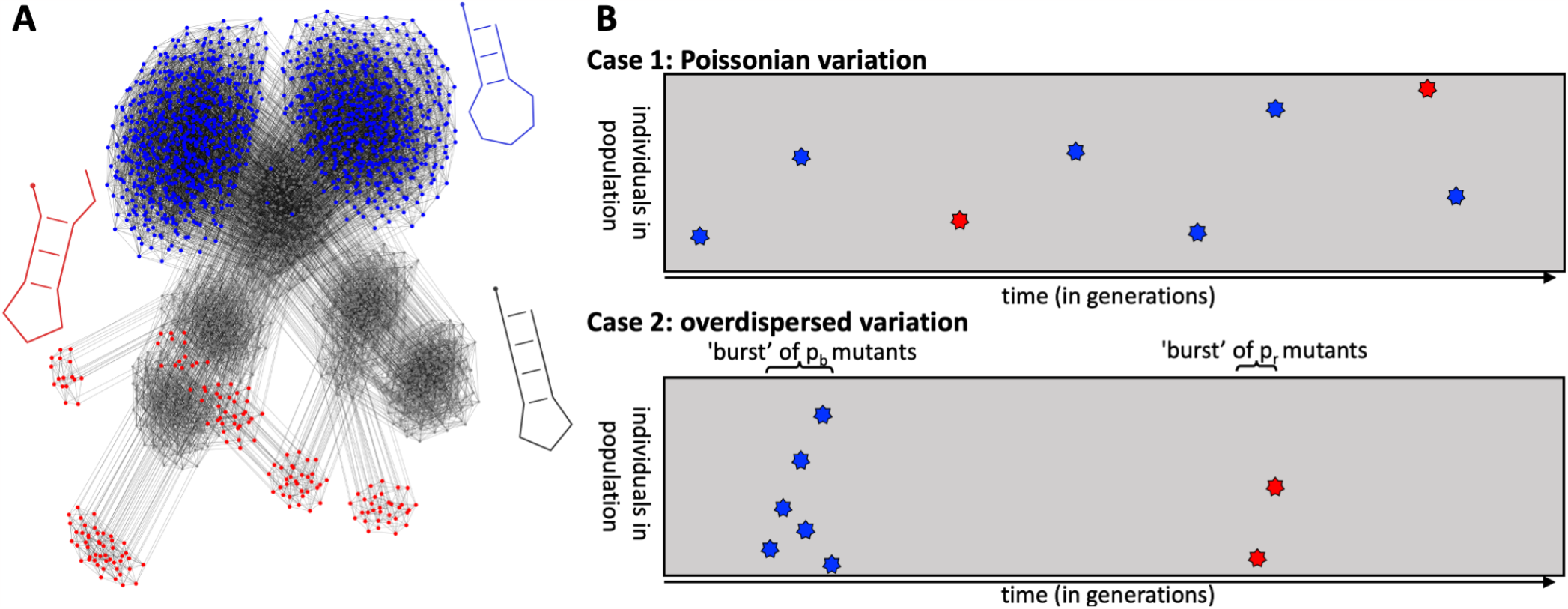
Structure in the mapping from genotypes to phenotypes can induce non-Poissonian bursts in the introduction of novel variation. **(A)** Genotypes mapping to three selected *L* = 12 RNA secondary structure phenotypes (shown in grey, red and blue) are drawn as a mutational network. Each genotype is a network node and each grey edge between two genotypes means that these two genotypes are only one point mutation apart. The graph is drawn using the force-layout algorithm from NetworkX [4]. The full neutral component (NC) of 1094 genotypes (nodes) is shown for the grey phenotype (specifically the NC containing the sequence AUACGAAACGUA), while only those nodes connected to the grey one are shown for the red and blue phenotypes. This network is heterogeneous in several ways: first, not all grey genotypes are portal genotypes with mutational connections to red or blue phenotypes. Secondly, the grey phenotype exhibits a modular community structure where the nodes form several clusters which are densely connected internally, but only have a few connections to other clusters. Thirdly, the portal genotypes to the red or blue phenotype are concentrated on a few regions of the grey network, i.e. transitions to blue or red are very likely from some grey genotypes and their mutational neighbours, but impossible otherwise. **(B)**Schematic plot of individuals in the population (y-axis) vs. time (x-axis). The population starts on the grey phenotype and moves through the grey NC by neutral mutations. Other novel phenotypes can appear through random mutations, but in this simplified case of strong stabilizing selection the novel phenotypes only appear for one generation. Here only two novel phenotypes, blue and red, are depicted, with blue appearing at a larger rate than red. Case 1 depicts the classical picture with Poisson statistics. Case 2 exhibits “bursty/overdispersed variation” due to the discrete nature of the genotypes and by heterogeneous structure in the GP map. Both cases have the same average rates of introduction. Note that each colour stands for one *phenotype*, but in this many-to-one mapping, this does not imply that they have the same *genotype*.

The simplest source of heterogeneity in the distribution of portal genotypes would be simple statistical fluctuations due to the fact that the probability 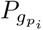 that a randomly chosen genotype is a portal to a specific phe-notype *p*_*i*_ is small, a condition thought to hold widely [7]. An example can be observed in Fig 1A, where many of the genotypes on the NC of the grey phenotype have neither a red nor the blue phenotype in their 1-mutational neighbours. Therefore, even if the portal genotypes were homogeneously distributed over the NC, we will observe bursts in the production of a specific phenotype *p*_*i*_ when a portal is present in a population, and no production of *p*_*i*_ otherwise (see the schematic in Fig. 1B) [31].

Moreover, Fig 1A depicts further inhomogeneities in the distribution of portal genotypes that go well beyond simple statistical number fluctuations, and which should lead to stronger non-Poissonian bursts. Indeed, we see a second type of heterogeneity: genotypes in the NC of an initial phenotype *p*_*g*_ typically form clusters (or communities from a network science perspective) where genotypes in the same cluster are more highly connected through mutations [11, 32]. Evolving populations can become trapped in a given part of the network for extended periods of time and only move into a new part rarely [33], in which case an average-rate model should be computed with community-dependent (and therefore varying) rates. A third and final potential source of heterogeneity we will class under the general heading of *non-neutral correlations*, namely that if a genotype is a portal for *p*_*i*_, then the probability of it having multiple mutations to *p*_*i*_ in its 1-mutation neighbourhood, or the probability that close neighbours are also portals for *p*_*i*_, can be significantly enhanced over a random model [22, 34]. All these sources of extra heterogeneity will strengthen the bursts. Indeed, some evidence of burstiness for specific NCs has been presented in ref [34].

In this paper, we will mainly explore the effect of such inhomogeneities with the RNA secondary structure GP map as an example. This is a suitable model because it is both biologically relevant for the function of non-coding RNAs and can be efficiently modelled with computational techniques using the ViennaRNA package [35]. While we focus mainly on the RNA GP map, we also repeat our analysis for two other GP maps, the HP model for protein tertiary structure [12], and the biomoprhs model of development [19]. Using these GP maps, as well as theoretical calculations, we systematically study the strength of the bursty dynamics, its dependence on population genetic parameters and its implications for adaptive evolution.

We proceed as follows: First, we build on simple scaling arguments from ref [31] to explore the time-scales and sizes of bursts, as well as their impact on adaptive dynamics for the simplest case of the fully monomorphic regime. Next, using a mixture of analytic and computational methods, we separate out the effects of different types of GP map inhomogeneities by constructing a hierarchy of null models. The simplest two models are the average-rate model and a random null model from [31] that has sequences randomly linked to phenotypes. The more complex models add increasing levels of non-random structure until the final level describes the full GP map. All levels of complexity are studied by population genetic simulations, and for the two simplest levels, we can also derive analytic descriptions of the statistics at which novel phenotypes appear through mutations. We repeat the simulations for a range of population sizes and mutation rates and find that, as expected [31], the introduction of new phenotypes is most strongly overdispersed for large population sizes and low mutation rates. Next, we study how bursts affect adaptive evolution in a landscape where one of the non-neutral variants has a selective advantage over the initial phenotype. We show that bursty dynamics can strongly increase average fixation times compared to an average-rate model. Moreover, the fixation rates saturate at a modest fitness threshold, and only weakly increase with fitness above this threshold. The root cause of these effects is that, with bursts, the discovery of a portal genotype is the rate-limiting step in the adaptive dynamics. Finally, we study the *arrival of the frequent* [31] (or *first come, first served* [3]) scenario for a two-peaked fitness landscape, where the fitter phenotype has a much lower average rate of appearance. We show that the probability that the fitter, but less frequent, phenotype, fixes first can be markedly suppressed compared to the predictions of average-rate models such as those used in refs [3, 36, 37], and argue that these effects can extend to more complex fitness landscapes.

### I. SCALING ARGUMENTS FOR BURSTS IN THE MONOMORPHIC REGIME

In this section we explore some simple scaling arguments in order to provide intuition for how inhomogeneities in the distribution of portal genotypes, sequences with at least one mutation to a desired phenotype, affect evolutionary dynamics. Consider a system with genotypes that are sequences of length *L* with alphabet size *K* so that the number of 1-mutational neighbours is *L*(*K−*1) which grows linearly with *L*. Note that the word genotype is typical of the GP map literature, but in practice this concept can stand for a gene, or any other unit of replication. The number of possible sequences grows exponentially with length as *K*^*L*^, which can quickly become hyper-astronomically large [38]. How the number of phenotype grows with increasing *L* is more complex, but it is reasonable to expect that it also grows exponentially with *L*, so that for large enough *L* any given sequence can have only a small fraction of all phenotypes in its 1-mutational neighbourhood. While these arguments need to be justified in more detail for any specific biological system, overall they imply that we should expect that a relatively small proportion of the set of all *K*^*L*^ possible sequences will be a portal genotype to any particular phenotype *p*_*i*_.

Here, we focus on the mutational introduction of a novel phenotype *p*_*i*_ in a population that is initialised on an initial NC of a phenotype *p*_*g*_ and initially evolves neutrally from genotype to genotype in this NC due to genetic drift. We mainly limit ourselves to the *weak-mutation* or *monomorphic regime*, where scaling arguments are relatively straightforward to make. We start by building on earlier work by McCandlish [34] and our group [31], where some of the arguments below were, to our knowledge, first mentioned. In the monomorphic regime, the product of the point mutation rate *u*, sequence length *L* and population size *N* is small (*NuL <* 1), i.e. only a small number of new mutations occur in the population in any given generation. Thus, the population will be localised on a single genotype *g*_0_ until the next neutral fixation, with an average time-scale of *t*_*ne*_ given by [31]:

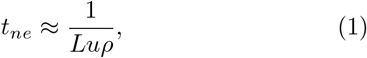

where the robustness *ρ* is the mean probability that a mutation is neutral on that NC. This time-scale is independent of population size *N*, because, as famously shown by King and Jukes [39] and Kimura [20], the fixation probability scales as 1*/N* which cancels out the *N* in the rate of new neutral mutations.

Similarly, the time-scale *t*_*gene*_ on which each specific mutation in the 1-mutational neighbourhood of a genotype of length *L* is produced is given by [31]:

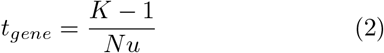

If the population is fixed on a genotype *g*_0_, then the mean number of times any specific mutation in the 1-mutational neighbourhood of that genotype will be produced before the population neutrally drifts to a new genotype is therefore given by the ratio [31]:

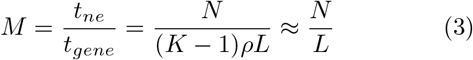

which is independent of the mutation rate. The final simplification follows because typically (*K−*1) is 3 (for DNA) and 19 (for proteins) while *ρ* is typically not too small [22] so that their product is typically a number roughly of order 1. We will call *M* the burst size since it is the number of times the same new genotype (and thus the same new phenotype *p*_*i*_) is introduced while the population is on a portal genotype for that phenotype *p*_*i*_. The true burst size will be larger if there is more than one mutation to *p*_*i*_ in the 1-mutational neighbourhood of *g*_0_, for example due to the inhomogeneities present in the RNA map. The time between such bursts of size *M* is set by the time-scale for the population to drift onto a portal genotype. If, as argued above, the probability 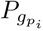 that a given genotype is a portal genotype to *p*_*i*_ is small, then the average time-scale to encounter such a portal genotype will scale as:

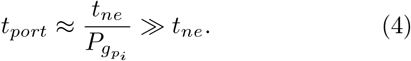

To summarise, in the monomorphic regime, if 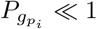 and *N/L≫*1, there will be long periods with no mutations to *p*_*i*_ until the population drifts onto a portal genotype, with a time-scale *t*_*port*_. If this portal genotype has 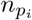 mutations to *p*_*i*_ in its 1-mutational neighbourhood, then (if *p*_*i*_ does not fix) the population will produce *p*_*i*_ an average of 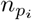*M* times before the population neutrally fixes to a new genotype on a time-scale *t*_*ne*_*≪t*_*port*_. Such a “burst” is illustrated in the second panel of Fig 1B. Since the appearance of the new phenotype *p*_*i*_ depends on a rare event (the fixation of a portal genotype), these appearances will be overdispersed, similarly to the case of noise in gene expression, where a small number of mRNA in a cell may produce a larger number of proteins in bursts [40].

Perhaps the most interesting impact of bursts is on the dynamics of adaptation: Consider a phenotype *p*_*i*_ with a single-mutant fixation probability 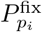. Now, if a portal genotype is found, on average a burst of *M* mutants of phenotype *p*_*i*_ will be produced. Then the probability 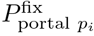 that a fixation event occurs by the end of that burst is well approximated by the following expression:

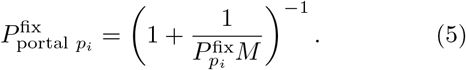

which is derived in the Appendix. When 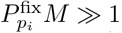, this function saturates towards 1 (see Appendix Fig 8), and its value is insensitive to changes in the single-mutant fixation probability 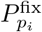, which depends on the selective advantage. In other words, if 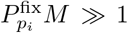, many more mutations that create a *p*_*i*_ occur than are strictly needed for fixation, so compared to an average-rate model, the fixation times can be substantially longer, as illustrated in Fig. 2. In other words, when the single-mutant-fixation probability 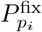 exceeds a threshold of 1*/M*, the time to fixation is primarily set by the timing of the first burst rather than by the strength of selection. Since for many models, 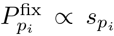 as long as 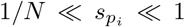, where 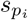 is the selection coefficient for *p*_*i*_, the saturation effect becomes relevant when

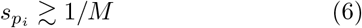

**FIG. 2.**
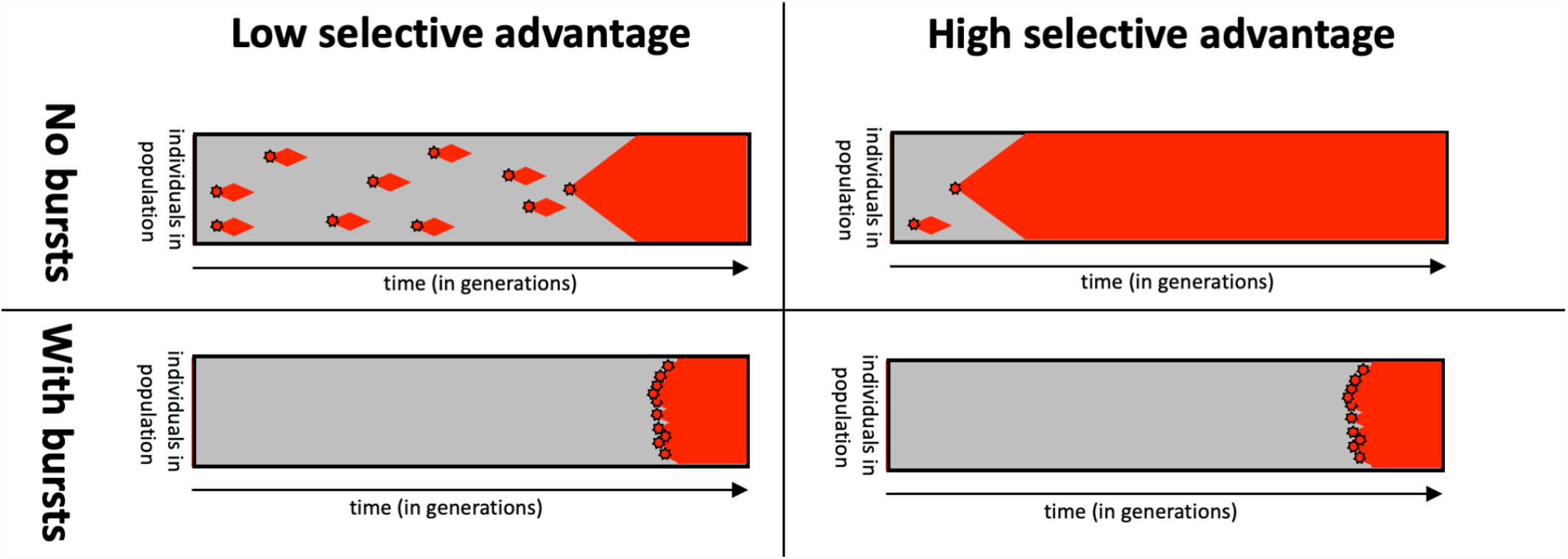
Idealised schematic - the effect of bursts on the time to fixation: in the non-bursty case (first row), red phenotype *p*_*r*_, which is fitter than the rest of the population, appears at intervals that are described by a Poisson distribution. The fixation time depends strongly on how many appearances of *p*_*r*_ are required for its fixation, which in turn depends strongly on the selection coefficient 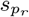 . In the overdispersed case (second row), there are time intervals where *p*_*r*_ does not appear at all for many generations and intervals when the population reaches a portal genotype, and *p*_*r*_ is produced many times in quick succession relative to the time-scale of neutral fixations. When *p*_*r*_ does not appear at all, it cannot fix, so its selective advantage does not matter. When it appears repeatedly, it may appear so frequently that it is likely to fix as long as its fitness is above a modest threshold given by Eq 6, how far above the threshold does not matter much. The time to fixation in this regime is dominated by the time *t*_*port*_ for the population to fix on a portal genotype. The fitness plays a much less important role.

If bursts are only caused by the simple random fluctuations captured by our scaling arguments, the threshold scales as *s*_*p*_ *> L/N* (see Eqns 3 and 6)^1^, but the threshold can be even smaller if other sources of overdispersion lead to larger burst sizes *M* . For simplicity, we have so far worked with an idealised monomorphic population that is always located at one genotype at a time. A fuller treatment of a more realistic monomorphic population is presented in the Appendix (section IV) and the resulting predictions are shown alongside our simulation data as cyan lines in Figs 4, 6 & 7.

**FIG. 3.**
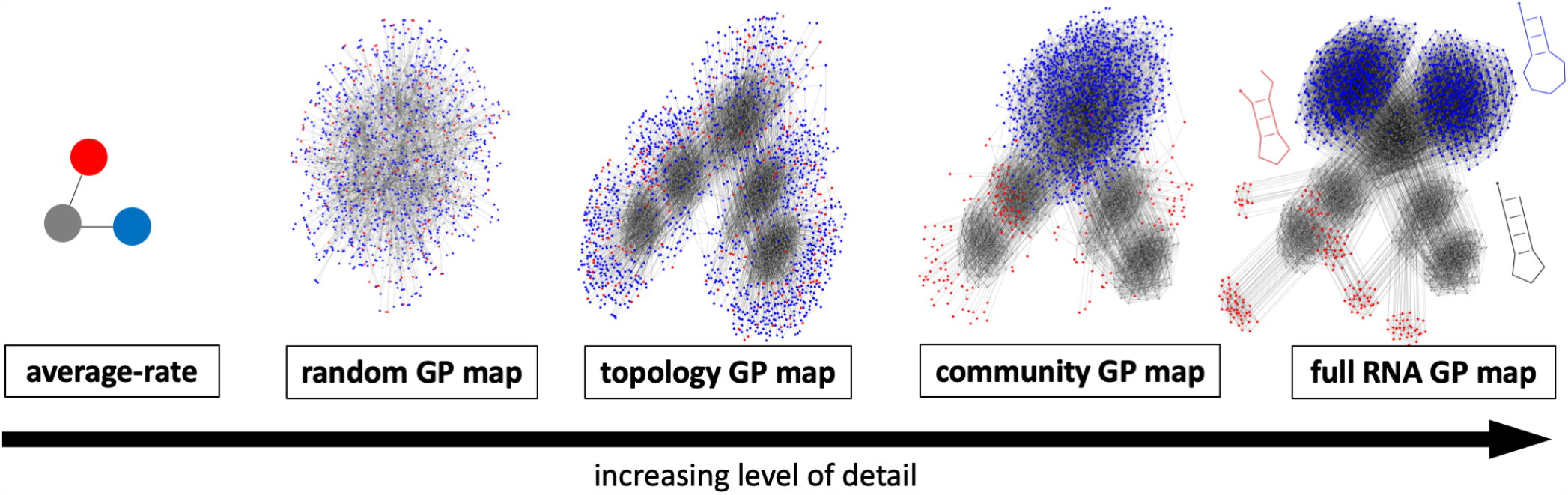
Hierarchy of models with increasing complexity for the RNA GP map. The rightmost network is the same as Fig 1: The 1094 genotypes in the initial NC, which corresponds to phenotype *p*_*g*_, are drawn as grey nodes, and possible point mutation connections are shown as grey lines. In addition to neutral mutations within the NC, mutations to two different non-neutral phenotypes are shown, 1358 genotypes with phenotype *p*_*b*_ (blue nodes) and 176 genotypes with phenotype *p*_*r*_ (red nodes). The leftmost model depicts a simple *average-rate model* without the internal structure of a GP map, but the same mean probabilities of mutating to *p*_*b*_ and *p*_*r*_. In the *random GP map*, the probability that a mutation from grey will lead to *p*_*i*_ is the same as in the RNA GP map, but otherwise the assignment between genotypes and phenotypes is random. The *topology GP map* has all the neutral connections of the original NC, but randomised non-neutral mutational neighbourhoods, thus erasing non-neutral correlations. The *community GP map* also randomises non-neutral mutational neighbourhoods, but only performs swaps within a network community, thus only partially erasing non-neutral correlations. The rightmost drawing represents the full NC from the *RNA GP map* and its mutational neighbourhood. The three structures are shown next to the full RNA map, as in Fig 1. To make the figure easier to interpret, only an excerpt is shown for the random GP map.

**FIG. 4.**
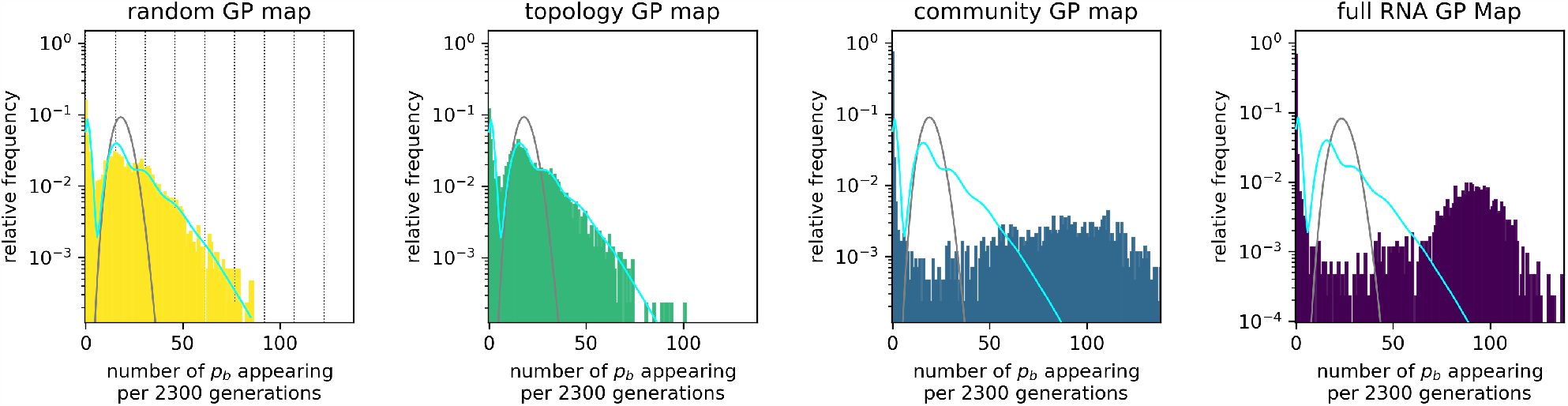
Strong deviations from Poisson statistics for the appearance of phenotype p_b_ in a population neutrally evolving with stabilising selection for p_g_. Phenotype mutation rates are measured by splitting the simulation into time intervals of Δ*t* = 2300 generations and recording how often the given new phenotype *p*_*b*_ appears in each Δ*t*. This data is shown for all four GP map models. The number of appearances per interval is highly overdispersed compared to a Poisson distribution with the same mean (grey line), which would be observed in the average-rate model. For the random GP map, the data can be approximated analytically (cyan line, given by Eq 17 in the Appendix). Vertical lines highlight the values 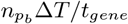 for a range of values 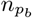, depicting the expected number of *p*_*b*_ mutants if a perfectly monomorphic population was located at a genotype with exactly 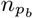 discrete instances of *p*_*b*_ in its mutational neighbourhood. The data on the community GP map and full RNA GP map shows even higher overdispersion than analytically predicted for the random map. Parameters: population size *N* = 1000, mutation rate *u* = 2 *×*10^*−*5^, total time 10^7^ generations. The initial NC is the one shown in Fig 3 and *p*_*b*_ corresponds to the blue phenotype in the same figure. Many further examples for other phenotypes, RNA length *L* = 30, and the HP and Biomorph GP maps can be found in sections 1.1 & 1.3 of the SI.

**FIG. 5.**
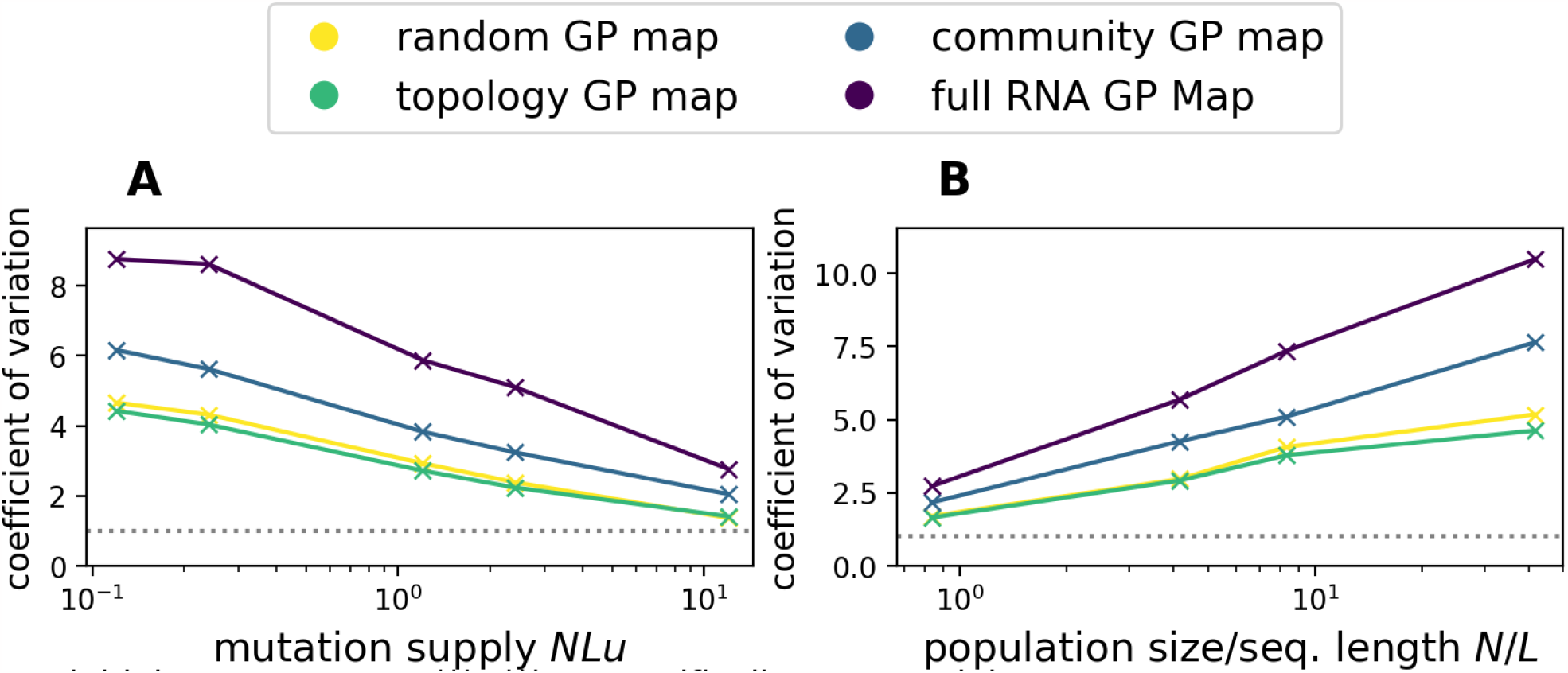
How does the amount of overdispersion, quantified by the coefficient of variation from Eq 7, depend on being in the monomorphic regime NuL≪1 or on being in the regime N*/*L*≫*1? We repeat the simulations from Fig 4: In A) we vary the mutation rate *u* at a constant population size *N* = 200, to study the effect of leaving the monomorphic regime. In B) we vary the population size *N* at a constant mutation rate *u* = 5 10^*−*5^ to study the fast-drift regime. The initial and final phenotype are the grey and red phenotype in Fig 1. Each line in the plot stands for a different GP map. The grey dashed line denotes the Poisson statistics prediction *V*_*t*_ = 1. Since the number of (neutral and non-neutral) mutations per generation scales as *NLu*, we need longer run-times to obtain reliable statistics for lower values of *NLu* and thus run simulations for *T* = max(10^6^*/*(*NuL*), 10^4^) generations, always rounded up to the nearest power of ten.

**FIG. 6.**
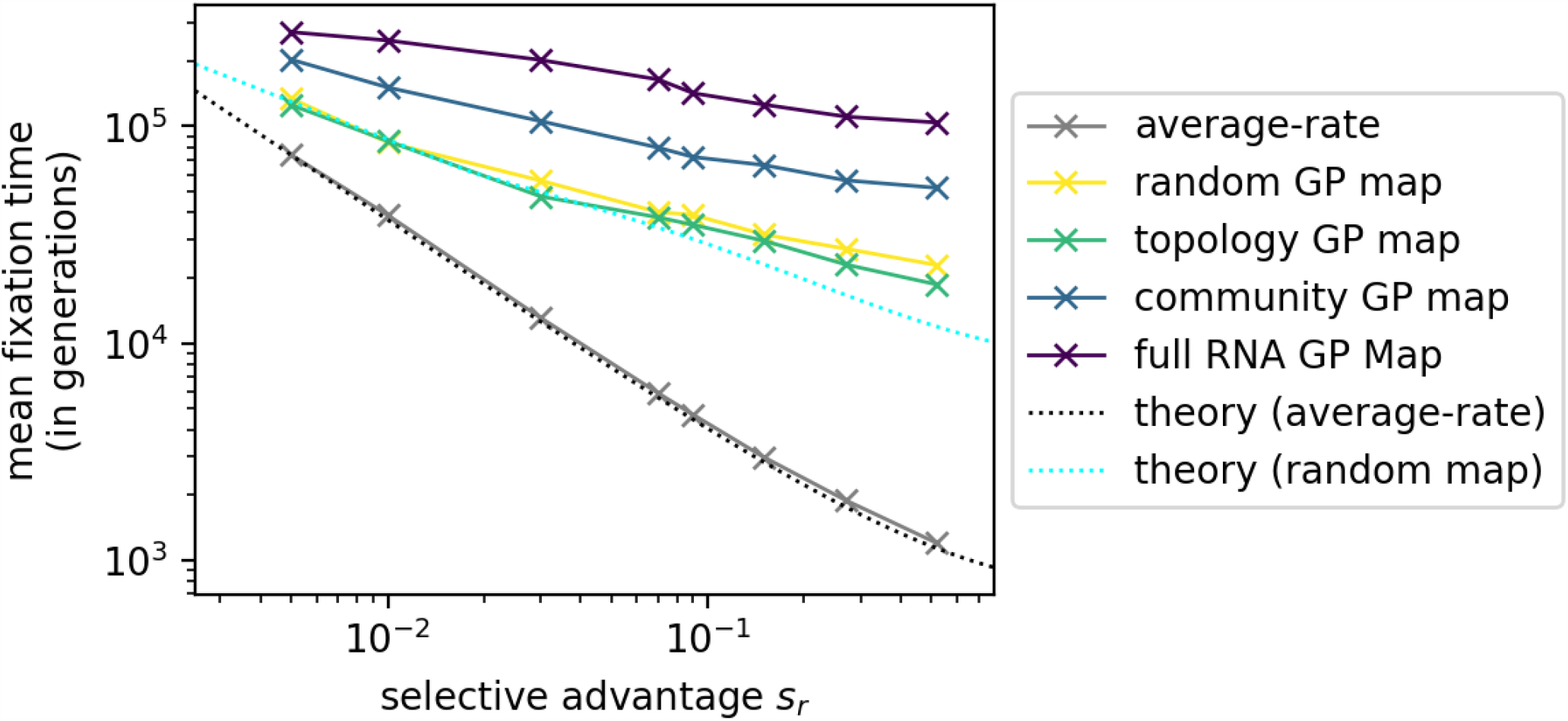
Overdispersion weakens the influence of changes in the selective advantage *s*_*r*_ of an adaptive phenotype on the time until its fixation. The population starts on the NC of the initial phenotype *p*_*g*_ and a single phenotype *p*_*r*_ has a selective advantage of *s*_*r*_ over *p*_*g*_ . All other phenotypic changes are deleterious with zero fitness. We repeat the simulation 10^3^ times for each value of *s*_*r*_ and record how many generations it takes on average from the start of the simulation until *p*_*r*_ fixes. Data is shown for all four model maps (the random, topology, community and RNA GP maps), as well as for an “average-rate” model, where variation is introduced by a random number generator at a fixed rate for each phenotype and a classic origin-fixation theory [2] (grey line, Eq 8) describes the data well. In all cases, *p*_*r*_ fixes more rapidly if its selective advantage is higher, but this decrease is much steeper for the average-rate model than on the GP map models which have overdispersed variation. The flatter scaling on the GP map models is captured by a simple analytic approximation for the random GP map (cyan dotted line, Eq 19). Parameters: population size *N* = 10^3^, mutation rate *u* = 2*×*10^*−*5^ so that *M≈*78 for the random map, and *NLu≈*0.24. The initial NC is the same as in the preceding Figs 3 - 5, and *p*_*r*_ is the phenotype drawn in red in Fig 3.

**FIG. 7.**
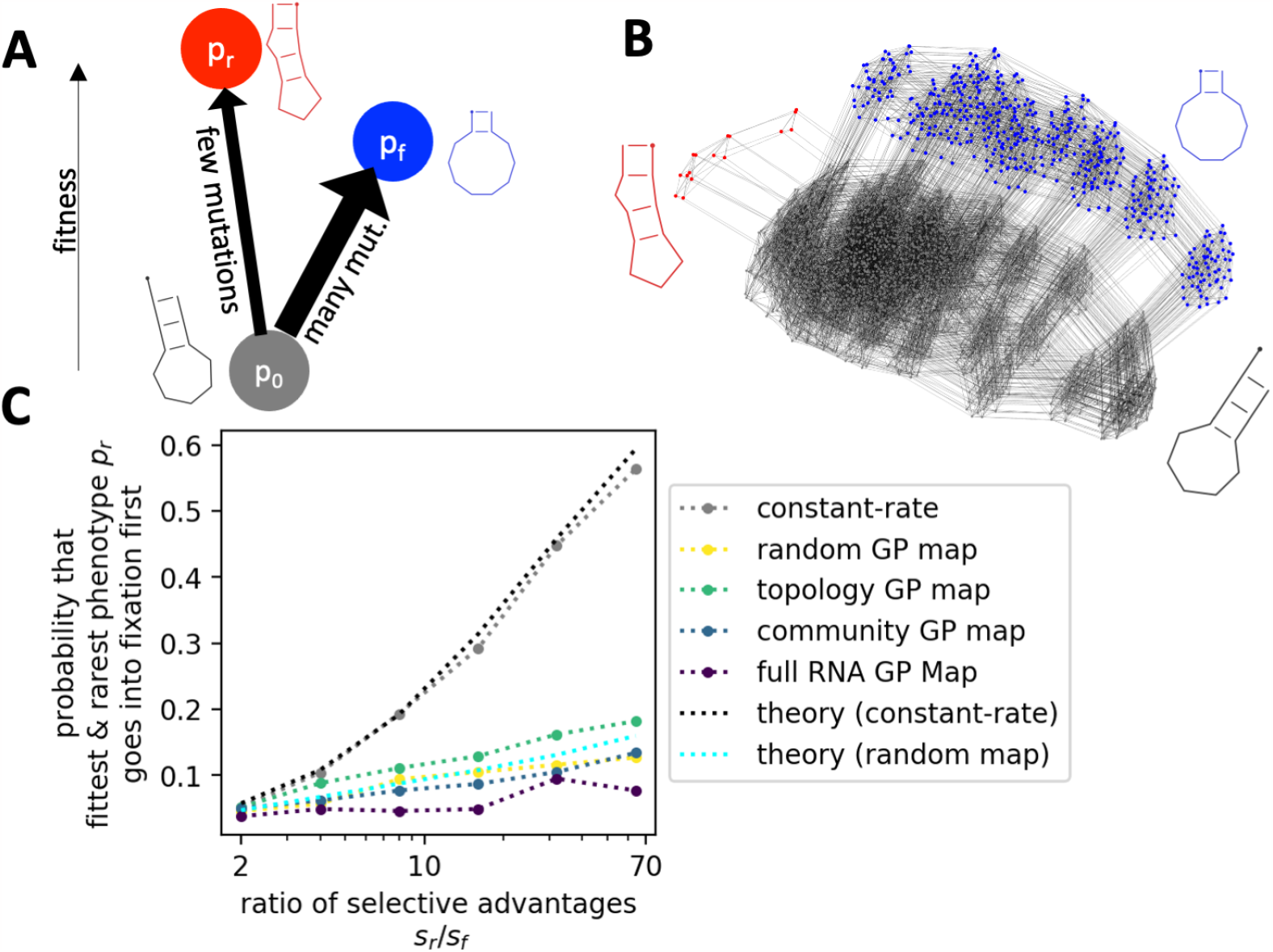
How overdispersion affects fixation probabilities for a landscape with two fitness maxima. A) Sketch of the fitness landscape (scenario from ref [31]) The population initially starts with phenotype *p*_0_ and can evolve towards one of two local maxima, phenotype *p*_*f*_ or *p*_*r*_. *p*_*r*_ is the global fitness maximum, but is less likely to arise through mutations (thus *r* for mutationally *rare* with 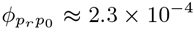 and *f* for *frequent* with 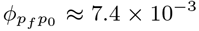). B) Sketch of the full mutational network relevant for this fitness landscape. C) For each of the GP map models from Fig 3, as well as for a case where *p*_*f*_ and *p*_*r*_ are introduced with constant rates, we record the probability that the fitter, but mutationally rarer phenotype *p*_*r*_ goes into fixation first. This probability is plotted against the selective bias towards *p*_*r*_, i.e. the ratio of the selective advantages *s*_*r*_ and *s*_*f*_, both relative to *p*_0_. In all cases, a higher selective bias towards *p*_*r*_ makes it more likely for *p*_*r*_ to fix, but this trend is less pronounced for the overdispersed case with the GP maps. The simulation results are well-predicted by theoretical calculations both for the random GP map (cyan dashed line, Eq 11) and for the average-rate model (grey dashed line, Eq 10). Parameters: *N* = 500, *u* = 2 *×* 10^*−*5^, probabilities based on 1000 repetitions.

By contrast, in the polymorphic regime *NLu≫*1, the population will carry a diverse number of genotypes at any time, and so inhomogeneities in the distribution of portal genotypes can be washed out [22, 31]. The dynamics and statistics will then be closer to that of an averagerate model. Nevertheless, as can be seen in Fig. 1A, the scale of the inhomogeneities in portal genotypes distributions across an NC can cover a significant range in Hamming distance. Therefore the strong inhomogeneities may cause bursty behaviour further into polymorphic regime than what would be the case for inhomogeneities that follow from statistical fluctuations due to small 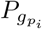 . Even at extremely high mutation rates, the inhomogeneity of the GP map can be important since high mutation rates typically entail a preference for high-robustness regions of a neutral set, which might be enriched in portals for some phenotypes over others [34].

Similarly, in what we will call the fast-drift limit *N/L <* 1, even monomorphic populations will produce new phenotypes with statistics more in line with an average-rate model [31]. The reason is that the population does not produce all genotypes in its one-mutational neighbourhood before moving on to a new 1-mutational neighbourhood through a neutral fixation. Again, structural inhomogeneities on larger Hamming-distance scales may still cause bursts in this fast-drift regime.

In the next section, we will use a combination of analytic and computational approaches to study in detail how the scaling arguments above apply to a specific system, namely the *L* = 12 RNA GP map.

### II. RESULTS FOR AN RNA GP MAP

#### A hierarchy of simplified models

In order to investigate how different features of the RNA GP map can lead to overdispersion in the arrival of novel phenotypic variation, we construct a hierarchy of simpler models that contain increasing amounts of the structure of the full RNA data. These are depicted in Fig 3, and discussed in more detail below.

At the first and simplest level, we use a model from ref [31], the *random GP map*, which has discrete genotypes, but no further correlations. In this map, the topology and genetic neutral and non-neutral correlations of the NC are completely erased by randomly assigning sequences of length *L* = 12 (the genotypes) to phenotypes (secondary structures) subject to the constraints that the mean outcomes of both neutral and non-neutral mutations are the same as the mean outcome of mutations on the grey NC in the RNA map. This map captures the fact that not every genotype on the NC is a portal genotype, just by simple statistical fluctuations.

At the second level, we define a *topology GP map*. Here, the initial NC and its internal topology (all neutral mutation connections) are identical to the full RNA map, but the phenotypic changes generated through non-neutral mutations are randomised (this approach is similar to a model found in ref [34]). The mean probability of a specific phenotypic change is set to match the corresponding NC in the RNA map, but *which* non-neutral mutation gives *which* phenotypic change is completely randomised. Thus, this map reproduces the neutral topology of the initial NC, capturing all neutral genetic correlations, but it will have no non-neutral genetic correlations.

At the third level we define a *community GP map*. As for the topology GP map, the initial NC and its topology are identical to the full RNA map. Unlike the topology GP map however, the randomisation of non-neutral mutations is applied to each network community of the NC separately. Thus, this map captures the fact that some phenotypic changes might be more likely in certain network communities of the RNA NC, but misses out on other kinds of non-neutral genetic correlations.

Finally, we investigate the full *RNA map*. In addition to the features present in the community GP map, we observe that non-neutral mutations to a specific phenotype are even more clustered to specific genotypes and their neighbourhood.

### Overdispersion in arrival rates on a GP map

We start with population dynamics simulations of the case where all non-neutral variants are unviable (i.e. have zero fitness). Although alternative phenotypes are introduced through mutations, they then disappear within one generation due to the strong stabilising selection. With this simplification, the population will be confined to the initial phenotype *p*_*g*_^2^ and we can study the introduction of new variation in isolation, without any ongoing non-neutral fixation processes.

To measure the statistics of the introduction of new mutations, we simulated a population of *N* = 1000 haploid individuals with a mutation rate per site of *u* = 2*×*10^*−*5^ using Wright-Fisher dynamics (see Methods). Since *NLu≈*0.24 this system is in the monomorphic regime. For fixed time intervals of Δ*t* = 2300 generations each, which is sufficient time for mutants to appear, but much shorter than the neutral fixation time scale of *t*_*ne*_*≈*12000 generations, we record how many times one specific new phenotype, *p*_*b*_, appears during each interval. We can easily detect overdispersion from this data: If phenotypic variation was approximately a constant-rate Poisson process, we would expect to see a relatively narrow distribution of *p*_*b*_ counts in each interval, with some fluctuation around the mean given by a Poisson distribution (grey curve in Fig. 4). However, as can be seen in Fig. 4, we see marked deviations from Poisson statistics for all four maps, and the deviation increases with map complexity. Intervals with zero appearances of *p*_*b*_ and intervals with a very high number of *p*_*b*_ appearances are much more common in the simulation data than for a Poisson distribution with the same mean. This is clearest for the full RNA map data, where only 0.4 % of all time intervals have *p*_*b*_ counts in the *μ±σ*(where *σ*is the standard deviation) range of the Poisson distribution: the counts in 73% of time intervals fall below this range (representing the time between bursts), while 27% appear above the range (the bursts). Similar findings hold for other phenotypes *p*_*i*_ (SI section 1.1), as well as for longer sequences of *L* = 30 (SI section 1.3), and for the GP maps of Richard Dawkins’ biomorphs and the HP protein model (SI section 2).

For the random GP map, the simplest of our GP map models, we can estimate the overdispersed distribution analytically, which provides a reasonably good fit to the data (cyan line in Fig 4). The full expression is derived in the Appendix (Eq 17) and captures the following simple phenomenology: if the population is perfectly monomorphic and remains on the same genotype throughout the time interval, then the expected number of *p*_*b*_ mutants produced is simply given by the number of *p*_*b*_ phenotypes in the mutational neighbourhood, 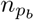, multiplied by the number of times each genotype in the mutational neighbourhood is expected to occur during Δ*T*, which is given by Δ*T/t*_*gene*_ (from Eq 2). During each Δ*T* the population is located at a genotype with an integer number of zero, one, two, etc… mutational neighbours that produce *p*_*b*_. Thus, we expect a peak at zero, and then successively smaller peaks at Δ*T/t*_*gene*_, at 2Δ*T/t*_*gene*_ etc. These values are shown as black dotted lines in the random map histogram, and match the peaks observed in the full distribution. The full analytic expression represented by the cyan line includes some further effects such as the fact that the population can fall off a portal genotype during Δ*T*, and that our simulations are not perfectly monomorphic, but the basic phenomenology is captured by the simple explanation above. This multi-peaked phenomenology is also clearly observed on data for another GP map, the protein HP model (Figs S14-S17 in the SI).

Having described the dynamics on the simple random map, let us compare all four GP maps. To help identify differences between the four distributions in Fig 4, the cyan line that approximates the distribution for the random GP map is included in all four subplots. First, note that the distributions from the random GP map and the topology GP map are quite similar. This similarity is perhaps not surprising, because in each map the portal genotypes are uniformly distributed across the NC. Next, we note that the overdispersion increases for the community GP map and even more for the full RNA GP map. These maps have an inhomogeneous distribution of portal genotypes over the NC. Thus, a population will not produce *p*_*b*_ when it is neutrally diffusing across areas of the NC that are depleted in portal genotypes for *p*_*b*_, and will repeatedly find portals when it is in a region that is enriched in them, leading to further overdispersion. The community structure of the neutral network may also imply that the time-scale to go from one part of the NC without portals to another part that has portals may be quite slow [34].

From these observations, we can deduce a number of factors that contribute to the overdispersion in phenotypic variation. First, the discreteness of mutations is a sufficient condition for overdispersion, as predicted by the scaling arguments in section I and shown here for the random GP map. Secondly, the similarity of the data from the random GP map and the topology GP map indicates that the topology of the NC in itself, which is caused by neutral correlations, may not lead to much additional overdispersion in the production of novel variation. Thirdly, the non-neutral genetic correlations that are present in the community GP map and the full RNA GP map, cause additional overdispersion. In the full RNA GP map, the distribution actually has a secondary peak at rates that are much higher than the mean. This extra peak is caused by the fact that it is no longer an exception to have several instances of *p*_*b*_ in a mutational neighbourhood since the few possible transitions to *p*_*b*_ are grouped around a very small part of the NC, as can be seen in Fig. 3. While the strength of these non-neutral correlations will depend on many details of the GP map [22] and differ for different target phenotypes [34], one simple source follows form the generic high robustness of all NCs [22, 41]: If a genotype that maps to the blue phenotype has several mutational neighbours that also map to the blue genotype, it is likely that a few of these are also mutationally accessible from one specific part of the grey NC.

### Influence of mutation rates and population sizes

So far, we demonstrated overdispersion in the arrival rates of non-neutral phenotypic variation for all four GP map models. However, we only performed this analysis for a single population size *N* and mutation rate *u*. Next, we investigate how these two parameters affect our results. For this analysis, we need a single number that summarises by how much the phenotypic variation found in a population deviates from a Poisson process. Here we consider the time intervals *t*_*r*_ between two successive and non-concurrent appearances of *p*_*r*_, which follow an exponential distribution in a Poisson process^3^, and focus on the coefficient of variation, defined as

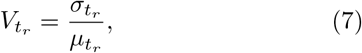

where 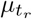 and 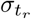 are the average and standard deviation of the time interval distribution. For a Poisson process, we would have 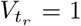 [42]. Higher values of the coefficient of variation indicate an overdispersed scenario, as in Fig 1B, where very short times between two *p*_*r*_ appearances (within ‘bursts’) and very long times between *p*_*r*_ appearances (between ‘bursts’) are common.

We simulate Wright-Fisher dynamics on all four maps for a range of mutation rates *u* and population sizes *N* and summarise the statistics by the coefficient of variation from Eq 7 in Fig 5. We can draw the following conclusions from the coefficients of variation: First, in agreement with the previous section, the community and RNA GP maps display the most overdispersed dynamics, and the random GP map and in the topology GP map are approximately similar.

Secondly, as can be seen in Fig 5A, overdispersion is strongest when the population is in the monomorphic regime, and 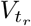 reduces as the population becomes more polymorphic with increasing *NLu*. The highest mutation rate in our data gives a highly polymorphic population with *NuL* = 12, where any two individuals are expected to have incurred≈24 mutations since their last common ancestor *N* generations ago. For this population, the random and topology map data are near the Poisson statistics expectation of 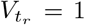, but the dynamics are still overdispersed for the full RNA GP map, suggesting that the population needs to be even more polymorphic before it spreads enough over the NC to wash out the larger-scale fluctuations in the distribution of portal genotypes illustrated in Fig. 1A.

Finally, as shown in Fig 5B, the overdispersion is weakest in the fast-drift limit where *N/L <* 1, since the population will move to a new neutral phenotype before the genotypes in its 1-mutational neighbourhood appear repeatedly to produce a burst. However, as before in the polymorphic regime, the dynamics on the full RNA GP map remains overdispersed even for small population sizes and thus strong drift, when other sources of inhomogeneity have washed out.

To sum up, we find that overdispersion is strongest when mutation rates are low and population sizes large as expected from the simple scaling arguments reviewed in section I. Moreover, for the full RNA GP map, we observe overdispersion in all cases, even though our data includes polymorphic populations with *NLu*≫1 and the fast-drift limit with *N/L <* 1, where the simple scaling arguments suggest bursts should disappear. These observations generalise to further phenotypes (see Supplementary Materials section 1.2).

### How bursts affect fixation times

In this section, we will test our earlier scaling arguments about fixation from section I. For simplicity, we consider a simple adaptive scenario, where only a single phenotype *p*_*r*_ has a selective advantage over the initial phenotype *p*_*g*_ and all remaining phenotypes are unviable. In this case the outcome is clear: at some point the fitter phenotype *p*_*r*_ will go into fixation. Nevertheless, exactly how many generations it takes for this fixation to happen will depend on the timing of the introduction of a new phenotype, and the strength of its selective advantage. The higher the selective advantage, the more likely an individual *p*_*r*_ mutant is to go into fixation and so the fewer *p*_*r*_ mutants are required for fixation and the lower the fixation time. As shown in Fig 6, this negative correlation between fixation time and selective advantage is indeed observed in all four maps, as well as for an average-rate scenario. However, the decrease of fixation time is much greater in the average-rate case than in the GP maps.

The basic reason for this weak dependence on fitness is described in section I. Once a selection coefficient is larger than a threshold that scales as 1/(burst size) the new phenotype will fix almost certainly during the first burst and so increasing the fitness further will only weakly affect fixation rates which are now set primarily by the arrival times of portals *t*_*port*_. These times are typically much longer than the average fixation time in the average rate model.

To test our quantitative understanding of the dynamics, we also include analytic approximations for the average-rate model and the random map in Fig 6. For the average-rate case, the time to fixation of *p*_*r*_ can be approximated as follows: in a classic origin-fixation model [2], we expect the mean fixation time *t*_*fix*_ to scale inversely with the product of an average-rate origin term *r*_*r*_ and single-mutant fixation probability 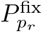(details in Appendix section IV):

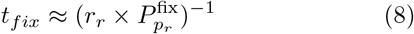

To match the GP map averages, the average-rate origin term should be 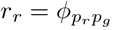 *NuL* where 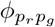 is defined as the probability that a phenotype *p*_*r*_ appears upon random mutations to genotypes that map to *p*_*g*_, averaged over the whole NC [31]. In other words, it is simply the mutation supply times the mean probability that a mutation generates *p*_*r*_. The second term, the single-mutant fixation probability for Wright Fisher dynamics, was calculated by Kimura [43] in the diffusion approximation, and for a haploid population takes the form [44]:

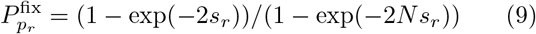

which only depends on the population size *N* and selective advantage *s*_*r*_ of *p*_*r*_, and is a reasonable approximation for a wide range of *s*_*r*_ [45]. The theoretical curve from Eq 8 and 9 is shown as a grey line in Fig 6, and approximates the numerical results from the average-rate model well.

For the random map, estimating the time to fixation of *p*_*r*_ is more complex and thus derived in the Appendix (section IV). The calculations (Eq 19) capture the basic scaling arguments from section I, but include some further effects that are relevant for our simulations, such as the presence of some neutral genotypic variation in the populations, which leads to small deviations from ‘perfect’ bursts described by the scaling arguments. The prediction is drawn as a teal line in Fig 6, and fits the data well for the random GP map and for the topology GP map.

To sum up, both our analytic calculations and simulation results indicate that the selective advantage of the adaptive phenotype *p*_*r*_ has a (much) lower impact on its fixation time in the overdispersed scenario than in the average-rate model, as was predicted in Section I.

### Implications of overdispersion for adaptation with two fitness maxima

In the previous section, the final outcome was always clear. There was exactly one phenotype with a selective advantage and this phenotype went into fixation in all simulations. The only question was its timing. In this section, we investigate a more general case treated for example in refs [3, 31], where two phenotypes, *p*_*f*_ (for ‘frequent’) and *p*_*r*_ (for ‘rare’), have selective advantages over the initial phenotype *p*_0_ and either of them could go into fixation first. These two phenotypes, *p*_*f*_ and *p*_*r*_, have different mean likelihoods to appear through random mutations, 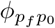 and 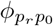, and they have different selective advantages, *s*_*f*_ and *s*_*r*_, over the initial phenotype *p*_0_, as sketched in Fig 7A. We are primarily interested in whether *p*_*f*_ or *p*_*r*_ will go into fixation first, as in refs [3, 31]. Therefore, we chose phenotypes *p*_*f*_ and *p*_*r*_ that are not connected by point mutations, such that both phenotypes constitute a local maximum that is difficult to escape from.

The most interesting scenario is when the less frequent phenotype *p*_*r*_ has the higher selective advantage, such that the bias in variation and selection favour different phenotypes. For the average-rate model, the probability *P*_r fixes_ that the fittest phenotype, *p*_*r*_ is the first to fix is given by a very simple ratio in the origin-fixation regime (derived in section IV in the Appendix, equivalent to the classic result from [3]):

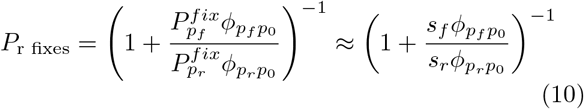

where the second approximate step holds for 1*/N*≪ *s*_*i*_≪1. There is only one effective parameter that sets the probability of the final outcomes, namely the ratio of the two origin-fixation terms. This simple analytic prediction is shown as a grey dotted line in Fig 7C and is in good agreement with our simulation results for the average-rate scenario. In other words, if we replace the GP structure of Fig 7B with average rates, then Eq 10 works very well.

How do the four different levels of GP map structure affect the probability of different outcomes? We observe in Fig 7C that a higher selective advantage of *p*_*r*_ still raises the probability that *p*_*r*_ fixes. However, the influence of this relative selective advantage is dramatically weaker for the bursty dynamics on the GP maps. The reason follows from our arguments about fixation times: Since both *p*_*f*_ and *p*_*r*_ have a selective advantage over the original phenotype, either of them is likely to fix immediately if it appears in a large burst. Then, as long as we are in the saturating regime of large bursts with *sM*≫1, the outcome is set primarily by whether the first burst to appear is one of *p*_*f*_ or *p*_*r*_ mutants, which in turn is set by probability of finding a portal genotype and not by selection. This intuition can be quantified if we assume that *p*_*r*_ and *p*_*f*_ appear and fix independently from one another. Then the probability that *p*_*r*_ fixes first depends on their individual fixation times 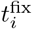 as follows (see Supplementary Information section 5.1):

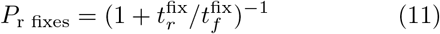

which reduces to the predictions Eq 10 if we use the average-rate expression (Eq 8) for the fixation times. Moreover, if we instead use the fixation times calculated using random GP map calculations, see Eq 19, and put them into Eq 11, then we find good agreement with the simulations for the random map (see the teal line in Fig. 7). Here, the reduced sensitivity to selective advantages in the fixation times also leads to reduced sensitiv-ity to selective advantages in the probability of a specific outcome. For the RNA map, the same qualitative arguments apply, but the time until a portal is reached and the burst size are set by a more complex distribution of portal genotypes, and thus cannot be as easily captured by a simple analytic expression. Given the success of Eq 11 for the random GP map, we show in SI section 5.2 that Eq 11 can easily be generalised to multiple peaks, as long as we assume that the different phenotypes are introduced independently from one another. If there are *n* phenotypes that may all fix, then the probability that phenotype *p*_1_, with fixation time 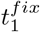 will fix before the others is given by:

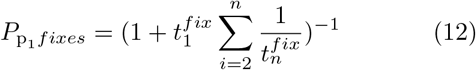

Although the analytic predictions for the two-peaked regime in Fig 7 use a more sophisticated fixation time estimation that accounts for mild polymorphism and finite burst sizes (eq 19), we will discuss the multi-peaked situation using simpler scaling arguments valid more deeply in the monomorphic regime: As long as the selection coefficients of a set of alternate potential phenotypes in a landscape are larger that 1/(average burst size), then mean fixation times *t*_*fix*_ should, to first order, be dominated by *t*_*port*_ for each phenotype in the monomorphic regime. Then the phenotype most likely to fix first will simply be the one with the highest accessibility of portal genotypes, regardless of relative fitness differences. Once the population fixes on a new phenotype, then both the relative selection coefficients and the set of accessible phenotypes with higher fitness will change. From this new set, it will again be the phenotype with the shortest *t*_*port*_ which is the most likely to fix, etc…. In this way, bursts can profoundly affect the manner in which a more complex fitness landscape is explored, something we will investigate in a follow-on study.

We demonstrated via explicitly simulations that in the presence of bursts, the first fixation event on the two-peaked landscape can depend much less on selective advantages of the peaks, and much more on how likely they are to appear as potential variation, than one would expect from average-rate models. We argue that the bones of this argument can be extended to multiple peaks, although of course the interaction with a full fitness landscape makes predictions more complex. Note that our scenario differs from the “survival of the flattest” effect [46, 47], which also predicts preferential fixation for phenotypes with larger neutral sets or higher robustness, but which only applies at high mutation rates. Similarly, arguments based on “free-fitness” [48, 49], can also be used to explain why phenotypes with larger neutral sets are more likely to fix. The free-fitness formalism is inspired by statistical mechanics, and depends on steady-state assumptions. It would be more appropriate for a fixed fitness landscape on much longer time scales, when the population has repeatedly transitioned between the different phenotypes. In this case, the details of the short-term dynamics would be less important, including the effects of bursts. Nevertheless, because phenotypes with larger neutral sets tend to have shorter *t*_*port*_ [31], all of these different limits above end up predicting a relative preference for phenotypes with larger neutral sets. The question of precisely where in biology we should expect each scenario to hold remains open.

## III. CONCLUSIONS AND DISCUSSION

### A. Main conclusions

Explicitly taking into account the finite length *L* of genotypes, sequences that represent biological units of replication, such as genes, leads to non-Poissonian bursts of size *M* in the arrival of novel phenotypic variation *p*, as long as some phenotypes are only accessible from a relatively number of ‘portal’ genotypes, and the population is in the weak-mutation or monomorphic regime [31, 34]. Here we explore these bursts in detail, showing in particular that they strongly affect adaptive dynamics.

To explore the effects of “bursty” statistics on the arrival and fixation of novel phenotypic variation, we focused on the RNA GP map. We measured the amount of overdispersion for a set of five models with increasing amounts of structure, which allowed us to separate out several different causes of overdispersion. Adding the full connectivity of the neutral component, e.g. including all neutral correlations, does not change the overdispersion by much beyond simply taking the discrete and finite nature of the genotypes into account. However, at the next two levels, the introduction of further non-neutral genetic correlations, such as those quantified by Greenbury et al. [22], can markedly increase the amount of overdispersion. Roughly speaking these non-neutral correlations imply that genotypes that have access to a novel phenotype in their one-mutational neighbourhood are clustered together in the NC, generating additional sources of bursty behaviour. We further demonstrated that, as predicted in ref [31], the amount of overdispersion in the introduction of a novel phenotype further depends on the details of the evolving population: genetically diverse populations or small populations undergoing rapid genetic drift average over several genotypes in the neutral component and therefore produce new phenotypes at a more constant rate than large monomorphic populations do.

By evolutionary dynamics simulations on the RNA GP map we gave explicit examples of where overdispersion in the arrival of novel phenotypic variation impacts the dynamics of adaptation. We showed how burst imply that fitness differences play a smaller role in determining fixation times and evolutionary outcomes than in a Poisson model with the same average mutation rates. This reduced influence of selective advantages can be understood from a simple argument: The number *x* of introductions of *p* that are needed for a successful fixation event strongly depends on the selective advantage. In average-rate models, the time until *x* mutants appear depends linearly on *x*, but in a bursty model, the time until *x* instances of *p* appear can be approximated by the time to the first burst, as long as the burst size *M > x*. Then the exact value of *x* and thus the selective advantage becomes less important. One consequence of these phenomena is that if there are multiple phenotypes that are fitter than the one on which the population resides, then bursts imply that the relative effect of differences in selection coefficients is much less important than it would be in an average-rate model. Instead, the probability of the population moving to a portal genotype, which depends on the frequency of the relevant phenotype as well as on the distribution of portal genotypes, plays a more important role in what eventually fixes. We explicitly demonstrated this effect for a two-peaked landscape, and show how to extend our analytic calculations to multiple peaks. the effect of overdispersion should also be important for evolutionary outcomes on more complex fitness landscapes. We hypothesize that the these effects of overdispersion may help explain why frequencies of phenotypes found in nature can in some cases (such as RNA secondary structures [50–52] and the topology of protein complexes [53]) follow biases in arrival rates over many orders of magnitude even when natural selection has also been at play.

### B. Dependence of overdispersion on the population genetic regime

Let us first review the conditions on the population parameters under which bursts are expected: Our analysis in Fig 5 indicates that bursts appear whenever the population is monomorphic (*NuL <* 1) and sufficiently large (*N/L >* 1), in agreement with the scaling arguments in section I and ref [31]. Burstiness persists for a larger parameter range if there are non-neutral correlations in the underlying GP map, such as in the community and full RNA GP maps. This is because populations need to spread out further over the neutral set to escape local heterogeneities, something they can achieve either through genetic diversity at high *NuL* or through fast genetic drift at low *N* . Thus, we observe burstiness on the RNA GP map even when *NuL≈a* or *N/L≈b* up to some finite constants *a >* 1 and *b >* 1 that depend on the strength of the non-neutral genetic correlations.

The second condition, that the population is sufficiently large for the bursts to appear (*N/L* ≳ *b*), is likely to be met in most realistic cases. The first condition however, that the population is monomorphic (*NuL* ≲ *a*), is more restrictive. For example, in the infinite-population limit commonly studied for GP maps [10, 23, 54] populations are highly polymorphic, and so are predicted to spread over very large numbers of distinct genotypes in a NC. This *NuL≫*1 limit is most likely to be appropriate for modelling microorganisms, which can have very high population sizes, or for RNA viruses, which can also have high mutation rates [28]. For vertebrates on the other hand, Lynch has estimated [55] that *Nu* is typically 0.00027 *< Nu <* 0.0010. In that case *NuL <* 1 for any genes with *L≤*1000. Of course the evolutionary dynamics for these classes of organisms are generally more complex than the simple model we used here, and so further work is needed to work out when and where the effect of bursts will be most prominent.

### C. Generalisation to other molecular and developmental phenotypes

Let us next turn to the conditions on GP maps for bursty dynamics: The minimum criterion is that only a fraction of genotypes in a neutral component are portals to phenotype *p*_*i*_. That this should be generically the case follows from fairly general scaling arguments, and also from the positive link between neutral set size and evolvability [21], where neutral exploration allows a larger number of novel phenotypes to be discovered than would be possible from a single genome. Another way of thinking about this aspect is in terms of epistasis since a low probability of a portal implies that the effect of a mutation depends on the genotype to which it is applied, even within a neutral set [56]. Not all kinds of epistasis would lead to bursts. Although epistasis is a necessary condition, it may not be sufficient as the follwing simple example shows: if every genotype has one mutation to *p*_*i*_, but this mutation is at different sites for different genotypes, this would not lead to bursts. Nevertheless, many GP maps exhibit epistasis in genotype-phenotype or genotype-fitness relationships [24, 57–60]. A more detailed investigation is needed to flesh out the links between epistasis, which is quite a broad concept, and the conditions for bursts, before drawing further conclusions. More direct evidence for the prerequisites for bursts come from GP map studies that have shown that different genotypes in a neutral set have different non-neutral mutational neighbourhood. Examples exist both in molecular GP maps (such as RNA [21, 61]) and higher-order GP maps (such as in a model of neural development [62] and gene regulatory networks [14]). Moreover, non-neutral correlations, i.e. cases where these differences exceed those expected in the random GP map, have been observed in a range of molecular GP maps (for example RNA, protein quaternary structure and protein tertiary structure [22]) and are likely to exist in more further GP maps. In fact, in Supplementary Information section 2, we show that both the HP model of protein folding and Richard Dawkins’ more macroscopic biomorphs model of development show pronounced overdispersion in the arrival of new variation. An important open question for the future is how strong these effects are for other GP maps.

One additional limitation of our results is that we restricted mutations to single nucleotide substitutions. However, since non-neutral correlations have also been found when single nucleotide insertions and deletions are included [63], our results should generalise to a broader range of mutations. This question also calls for further study.

### D. Overdispersion and punctuated equilibrium

The effects of overdispersion sketched in the schematics in Fig 1 & 2 is reminiscent of the concept of punctuated equilibrium, which postulates that evolution proceeds in long periods of neutral evolution interspersed by short periods of rapid phenotypic changes [64]. While the overdispersed variation analysed here can lead to such punctuated patterns, even an average-rate model would predict long periods of stasis until a rare phenotypic transitions *p*_*i*_ appears and goes into fixation [65]. In previous work showing punctuated dynamics on GP maps [66–68], both of these factors are likely to have played a role: both the overdispersed appearance of *p*_*i*_ and the long wait-times for rare *p*_*i*_. There are further potential causes of punctuated dynamics, for example fitness valleys [69], and therefore one should not invoke the overdispersion of new phenotypic introductions as an explanation for punctuated evolutionary dynamics without considering these alternative explanations.

### E. Future work

The strong effect of GP map structure on the statistics of the introduction of novel phenotypic variation observed here raises many directions for future research. Firstly, there is the question of the strength of non-neutral correlations that amplify bursts beyond the simpler arguments based on discrete genotypes. This can only be addressed with more detailed ways to quantify these correlations in different GP maps and by using the results in further calculations. There is a large parameter space to explore, with different GP maps, different NCs, and of course parameters such as population size and mutation rate.

Secondly, we derived expressions for the relative fixation rates for a two-peaked landscape, or a multi-peaked landscape with one key assumption: that the introduction processes of *different* phenotypes are independent from one another. This assumption would break down, for example, if the mutational connections to two or more phenotypes of interest were clustered around the same part of the NC. Future work should address such phenomena both analytically and computationally.

Thirdly, we have worked with a GP map, where each genotype corresponds to a single phenotype. Further questions arise around GP maps that have a non-deterministic relationship between genotypes and phenotypes [70]. Similarly, the concepts should be applied to transcription factor binding landscapes. Since *L* is typically short in this case, bursts could also play a role, but it is also a more complex GP map, where each genotype can bind to multiple transcription factors with a varying quantitative binding strength [71].

Furthermore, our analytic approximations only use an approximate treatment of mildly polymorphic populations. While this is sufficient to estimate slight deviations from a perfectly monomorphic population, future work should provide analytic approximations for populations on GP maps that fill the gap between the idealised cases of highly polymorphic populations, the infinite-population limit, and the weak-mutation-strong-selection/monomorphic limit.

Finally, the big question is how to observe these effects experimentally. Unlike previous work on burstiness on a genotypic level [25, 28], including interesting parallels to work on clusters of identical mutations originating from a premeiotic mutational events [72], our bursts are fundamentally a *phenotypic* effect that cannot be identified from sequences alone. There has been significant work showing that mutational biases can be observed in a wide range of organisms [73], but these effects are also seen at the sequence level. To illustrate, why bursts are a phenotypic effect, let us consider the schematic in Fig 1. On a genotypic level, we would see a different picture: first, there are neutral genetic changes that maintain the initial (grey) phenotype throughout the process. Secondly not all the blue mutants in the first burst have to have the same genotype, even though they all have the same phenotype since a portal may have several mutations to the blue phenotype due to non-neutral correlations. Thus, the bursts described in this paper can only be observed if we have both *genotypic* and *phenotypic* information for an evolving population for the entire population, not only mutations that are close to fixation, but also mutations that have just been introduced. Despite these difficulties, experimental tests of our predictions could be designed and performed.

## IV. METHODS

### RNA GP Map

For the RNA GP map, we folded all possible sequences of length *L* = 12 with the ViennaRNA package [35] (default parameters, version 2.4.14). We took each sequence’s folded structure as this genotype’s phenotype, but considered sequences to be non-folding if the mfe criterion was met by two degenerate structures. NCs are constructed in NetworkX [4] and drawn with its force-layout algorithm.

### Hierarchy of GP maps

We start by identifying all sequences that belong to a given neutral component (NC), i.e. a mutationally connected set of sequences folding into the same phenotype [24]. Then we choose one initial NC in the RNA GP map to build simpler models for this NC. First, we determine the mean mutation rates 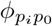for that NC, i.e. what fraction of mutations starting at this NC give a specific new phenotype *p*_*i*_. Note that all 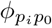 sum to one by definition when the probability of neutral mutations, the robustness 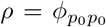, is included. We also identify the network communities of this NC following Weiß and Ahnert’s [11] method. Then the three simplified models were constructed from this NC information.

**Random GP map:** in this map, phenotypes are assigned to genotypes at random and the only input are the frequencies of each phenotype [31]. Here, we set the frequency of phenotype *p*_*i*_ to 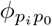, so that the mean mutation probabilities will match those for the initial NC in the RNA map.

**Topology GP map:** here the genotypes that form part of the initial NC are left unchanged, so that the topology of this NC matches the one in the RNA map. The unchanged NC topology already ensures that the fraction of neutral mutations matches that in the RNA map. All remaining genotypes are assigned random phenotypes (except the initial phenotype *p*_0_), each with a probability proportional to the rate from the RNA NC, 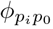. Here these probabilities had to be renormalised so all 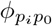 without the neutral mutations for *p*_0_ sum to one.

**Community GP map:** here, we start with the full RNA data and randomise the mutational neighbourhood of one community in the initial NC at a time: for each community we identify all genotypes that are mutational neighbours to this community, but not to another community in the initial NC. We shuffle the phenotypes associated with these genotypes to randomise the non-neutral mutations within each community. In order to leave the mean mutation probabilities intact, we identify subsets of genotypes that have exactly *n* connections to the NC, and only perform swaps within each subset.

### Simulations of evolving populations

For the evolutionary dynamics simulations on GP maps, we followed previous studies of evolutionary simulations on GP maps [31, 74] and implemented a Wright-Fisher model of a fixed number *N* haploid individuals in Python. Mutations were modelled to occur with constant probability *u* per reproduction event and site and the pheno-type of the mutated sequence was given by the GP map. The population was initialised on a single genotype in the selected NC, and then evolved neutrally for 10*N* generations before any data was collected, in order to randomise these forced initial conditions, as in [31]. We considered a fixation event to have occurred if less than 25% of the population carried the initial phenotype. In order to exclude rare and irreproducible jumps to other NCs of the neutral set of the initial phenotype *p*_0_, which would lead to non-homogeneity in the observed variation and thus confound our analysis, we set only genotypes in the initial NC of *p*_0_ to fitness 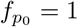.

In order to simulate the average-rate scenario, we also performed simpler simulations without the GP map. Here, we simply assumed that *L×u* mutations occur per individual and generation, in order to match the GP map case, where the mutation rate is given per site. In the average-rate model, each mutation has the same probability 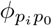 of giving phenotype *p*_*i*_ and a constant probability 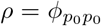 of leaving the initial structure unchanged. These rates are free parameters in the average-rate model, which we set to match the corresponding GP map values for the initial NC. For the rare event that a phenotype *p*_*j*_ different from the initial phenotype exists in the population and mutates, we simply set rates that match the mean rates for mutations on that phenotype in the RNA map (rather than a specific NC of that phenotype).

## APPENDIX: ANALYTIC CALCULATIONS

In this section, we will derive analytic expression for the expected rates, fixation times and fixation probabilities, first for the average-rate model from ref [31] and then for the overdispersed dynamics on the random GP map. In both cases, we can treat the system using a simple statistical treatment. If we wanted to model the full RNA map, we would need a more detailed treatment that explicitly includes the non-random, inhomogeneous structure of the GP map, such as the matrix treatment by McCandlish [34]. To simplify calculations, we ignore the time required for an ultimately successful mutant to actually go into fixation. All key variables for the calculations are summarised in table I.

**TABLE I.** Symbols used in the calculations.

| Symbol | Definition |
| --- | --- |
| $L$ | sequence length |
| $u$ | mutation rate (per site) |
| $N$ | population size |
| $s_i$ | selective advantage of phenotype $p_i$ |
| $\phi_{p_i p_0}$ | probability that a random mutation converts phenotype $p_0$ into $p_i$ (on the relevant NC), see ref [31] |
| $\rho$ | robustness, probability that a random mutation leaves phenotype $p_0$ unchanged (on the relevant NC), see ref [31] |
| $r_i = LNu \times \phi_{p_i p_0}$ | rate of $p_i$ introductions in the average-rate model [31] |
| $t_i^{\text{fix}}$ | expected time until $p_i$ is introduced and goes into fixation (derived for the average-rate case in Eq 13 and for the random GP map in Eq 19) |
| $t_{ne} = (uL\rho)^{-1}$ | mean time between neutral fixations through drift (see Eq 1 and ref [31]) |
| $t_{gene} = \frac{K-1}{Nu}$ | mean time until a specific mutation in the current mutational neighbourhood is observed (see Eq 2 and ref [31]) |
| $M = \frac{t_{ne}}{t_{gene}} = \frac{N}{(K-1)L\rho}$ | average burst size in the simplest approximation of the random map (perfectly monomorphic population, see Eq 3 and ref [31]) |
| $f_0$ | fraction of the population at the most prevalent genotype $g_0$ in our approximation of mildly polymorphic populations (derived in Eq 15) |
| $P_{p_1}^{\text{fix}} = \frac{1 - \exp(-2s_1)}{1 - \exp(-2Ns_1)}$ | single-mutant fixation probability for a single mutant of phenotype $p_1$ with selective advantage $s_1$ [44] (in the diffusion approximation) |
| $P_{\text{portal } p_1}^{\text{fix}}$ | probability that $p_1$ goes into fixation before the population leaves a given ‘portal’ genotype (derived in Eq 18) |

### Analytic calculations for the average-rate model

Modelling mutations to *p*_1_ as a simple Poisson process, i.e. occurring at a constant rate, as done e.g. in refs [31, 74], is the simplest approach. When selection also plays a role, we assume the origin-fixation regime, in which mutants are rare enough that each fixation process can be treated as an independent event and thus fixation probabilities can be written a simple product of mutation rates and fixation probabilities [2].

#### Poisson distribution of appearance events (grey line in Fig 4)

In the average-rate case, the number of *p*_1_ mutants per time interval is a Poisson distribution. This distribution only has a single free parameter, the mean. In Fig 4, we approximate the mean by the mean number of *p*_1_ mutants per time interval in the corresponding GP map simulation.

#### Fixation times as a function of s_r_ (grey line in Fig 6)

*p*_*r*_ mutants are introduced at rate 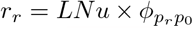 in the constant-rate model (ref [31] and table I) and the single-mutant-fixation probability is given by 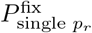 (see table I). Thus, if we ignore the time of the fixation process itself, and focus on the expected time until a *p*_*r*_ appears that will trigger a successful fixation event, we have:

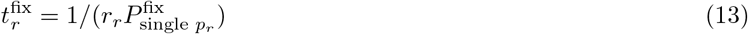

#### Fixation probability of two competing mutants (grey line in Fig 6)

In the origin-fixation regime, we can assume that all fixation events are independent and thus we model the fixation of *p*_*f*_ and *p*_*r*_ as independent processes occurring at constant rate with time scales 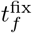 and 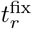, which are given by Eq 13. Then it can be shown (details in section 5.1 of the *SI*) that the probability that *p*_*r*_ fixes before *p*_*f*_ is given by 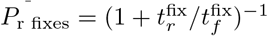.

In the limit *s*_*f*_ *≪* 1 & *s*_*r*_ *≪* 1 & *s*_*f*_ *N ≫* 1 & *s*_*r*_*N ≫* 1, we have 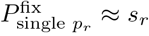, and so the expression can be rewritten as *P*_r fixes_ *≈* (1 + *vf*)^*−*1^, where *v* is the bias in variation, which is 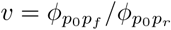, and *f* = *s*_*f*_ */s*_*r*_ the bias in selective advantages. This form agrees with Yampolski and Stoltzfus’ [3] well-known result (after renormalisation since they compute the ratio *P*_f fixes_*/P*_r fixes_).

### Analytic calculations for a simple overdispersed case: the random GP map

The key aspect to model in the random map is the following: a population can move to a new genotype through neutral drift without changing its phenotype, but the new genotype can have different phenotypes in its mutational neighbourhood. In the random GP map, such differences in the mutational neighbourhoods of different neutral genotypes are simple random fluctuations and can thus be modelled mathematically. Therefore, we can make analytic estimates for this case, relying on existing work in ref [31], where this model was first proposed.

As soon as we use a GP map, we need to account for the fact that a population of individuals with phenotype *p*_*g*_ can be made up of a mix of genotypes from the relevant NC of *p*_*g*_. To keep our calculations simple, we will assume that one genotype, which we will call the ‘prevalent genotype’ *g*_0_, dominates the population. In other words, we are treating cases close to the monomorphic limit. Then we will approximate the population dynamics based a single quantity, the prevalent genotype’s frequency in the population *f*_0_. In this approximation, the probability that two individuals in the population carry the same genotype, i.e. the homozygosity *H*, is well-approximated by the probability that two individuals carry *g*_0_, i.e. we have 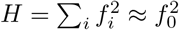(since the frequency of any other genotype *f*_*i*_ is low enough for 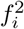 to be negligible).

We can thus estimate *f*_0_ if we know the homozygosity *H*, which we can approximate with coalescents: the probability that two individuals in a population of size *N* have a common ancestor in the previous generation is 1*/N* (1*−uL*(1*−ρ*)) (since the next generation is chosen from the *N* (1*−uL*(1*−ρ*)) individuals without deleterious mutations), and so on average the last ancestor is *N* (1*−uL*(1*−ρ*)) generations ago. Since the common ancestor, neutral mutations happen at rate 2*uLρ*. Using again the expression for two exponential processes (section 5.1 of the *SI*), the *homozygosity H*, the probability that no neutral mutation has taken place in either of the two lineages since the common ancestor is approximately:

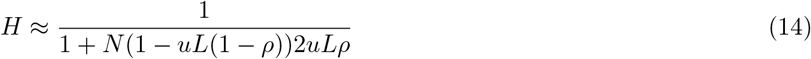

Using our previous expression 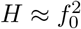, we can solve for the frequency of the dominant genotype *f*_0_:

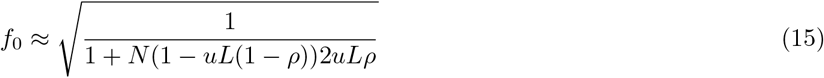

Note that we made a number of assumptions in this derivation, so it can only serve as a first approximation. The approximation is likely to get worse in the highly polymorphic limit (*NuL≫*1), but it has the correct limiting behaviour in the monomorphic limit (*f*_0_→1 as *NuL*→0) and it is found to be a sufficiently good approximation for the mildly polymorphic limit we encounter for the values of *u, L* and *N* that are used in this paper (see Supplementary Information section 4).

#### Distribution of appearance events over Δt (Fig 4)

In order to model, how many times we expect a phenotype *p*_*b*_ to appear through random mutations, we will need to model how many instances of *p*_*b*_ exist in the mutational neighbourhood of the current prevalent genotype. In the random GP map, differences between the mutational neighbourhoods of different genotypes are caused by simple statistical fluctuations around well-defined mean fraction of 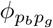 . Therefore, a binomial distribution can be used to describe the probability of finding 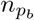 instances of *p*_*b*_ among the (*K−*1)*L* possible mutations in the mutational neighbourhood of a genotype *g*_0_:

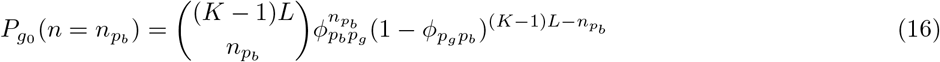

Note that, for computational efficiency, we used a phenotype with 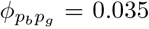 in Fig. 4. Then the probability of having portals with, for example 1, 2 or 3 1-mutational neighbours mapping to phenotype *p*_*b*_ is given by 0.36, 0.23, 0.09 respectively.

Next, we need to calculate how many *p*_*b*_ mutants we expect over a time interval Δ*t*, given that we start with a prevalent genotype *g*_0_ that has 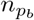 instances of *p*_*b*_ in its mutational neighbourhood. Here, we simply assume that *p*_*b*_ mutants arise at a constant rate over a time scale Δ*t* - but unlike the average-rate model, this rate itself depends on the current genotype and its value of 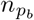 . However, our calculations also need to account for the fact that the dominant genotype may change during Δ*t*, and so we will sum over the probabilities of two cases:

- If the prevalent genotype in the population changes (i.e. a neutral fixation occurs) in the first half of the time interval, which occurs with *P* (neutral fixation middle) = 1*−*exp(*−*0.5 Δ*t/t*_*ne*_): then the population averages over the mutational neighbourhoods of two genotypes, and we simply approximate this scenario with the average-rate model (see table I), so we expect *r*_*b*_Δ*t* mutations of type *p*_*b*_.
- Otherwise, we assume that the prevalent genotype *g*_0_ is the same during Δ*t*: since each genotype in the mutational neighbourhood is produced every *t*_*gene*_ generations (Eq 2), and there are 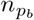genotypes with *p*_*b*_, *p*_*b*_ is produced at a rate 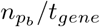. This rate applies only to the well-localised fraction *f*_0_ of the population; let us assume that the average-rate model provides a good approximation for the remainder of the population. Thus, in this scenario the number of *p*_*b*_ mutations expected in Δ*t* is 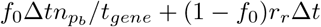.

Given the information on the rate at which we expect *p*_*b*_ to arise during a time interval Δ*t*, the probability of finding exactly *M* mutants can be calculated using a Poisson distribution Pois(*M, x*) with mean *x*. This gives:

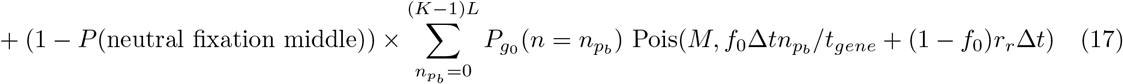

Here, the first term is reminiscent of the classic average-rate expression, which would be Pois(*M, r*_*b*_Δ*t*) and corresponds to cases, where genetic drift smooths over differences between genotypes in the NC. The second term gives rise to the overdispersed behaviour: essentially fluctuations in the number of *p*_*b*_ per mutational neighbourhood are described by the probability distribution 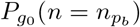 and these fluctuations then give rise to large rate variations in the Poisson distribution.

#### Fixation from a portal genotype

A ‘portal’ [30] genotype is any genotype with *p*_1_ in its mutational neighbourhood, i.e. 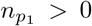. Let us assume that a population is currently located at a portal genotype, which has exactly one instance of *p*_1_ in its mutational neighbourhood (i.e. one out of 3*L* mutations from the portal genotype gives *p*_1_). Then the population is experiencing a burst until the portal genotype changes through genetic drift. Let us compute the probability that *p*_1_ will fix during that burst, i.e. before the portal genotype is lost through genetic drift on the neutral network. We can simply treat the two outcomes, a fixation to *p*_1_ or a neutral fixation on the network, as independent, exponentially distributed random processes and compute the likelihood that the *p*_1_ fixation occurs first. The timescales of these two random processes are:

- **Timescale of** *p*_1_ **fixation:** *p*_1_ appears every *t*_*gene*_ generations while the population is at the portal (see Eq 2). The probability that a particular *p*_1_ mutant goes into fixation is given by the single-mutant fixation probability 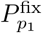 (table I). Then the time to *p*_1_ fixation is therefore approximately 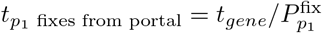.
- **Timescale of neutral fixation:** *t*_*ne*_ (see table I).

Outside the strict monomorphic regime, and under the assumption that the fitter phenotype is only accessible from the portal genotype, the calculation only applies to a fraction *f*_0_ of the population size, so we simply need to replace the population size *N* by *Nf*_0_.

Thus, we can use the expression for the ordering of two exponential random processes (SI section 5.1) to compute the probability that *p*_1_ fixes before a neutral fixation removes the portal genotype from the population:

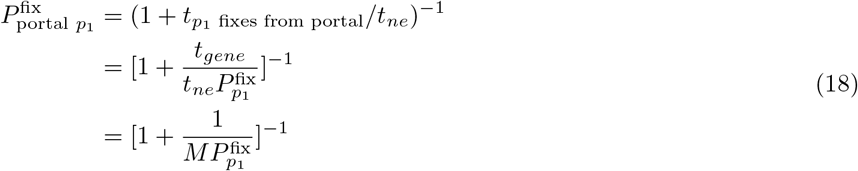

In the last line, we have written this expression in terms of the burst size *M* from Eq 3. This function is sketched in Fig 8 and we can distinguish two limiting cases:

**FIG. 8.**
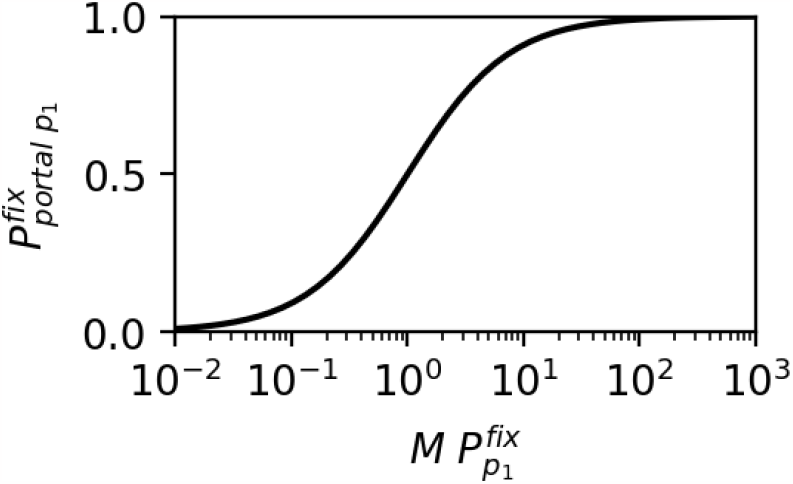
Probability of successful fixation event after a single burst on the random GP map 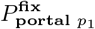, as a function of the typical burst size *M* and the single-mutant fixation probability 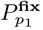: for large enough burst sizes *M*, the function has saturated and the probability of a successful fixation within that burst is close to one, regardless of the exact value of the product 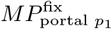. This means that the outcome is less sensitive to small changes in the single-mutant fixation probability 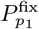, which in turn depends on the selective advantage.

1. If 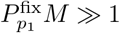, then we have 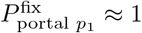. In this limit, a typical burst contains more than enough mutants for a successful fixation event. On the order of 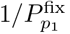 mutants are needed for the phenotype to have a 50% chance of fixing, so if *M* is much larger, then the probability of fixing is large, and will remain insensitive to small changes in the selective advantage, as long as the system remains in this regime. The time-scale for fixation will therefore be dominated by the time between bursts, rather than by 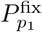.
2. If 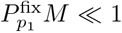, then 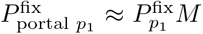. Then the fixation probability from a portal phenotype is proportional to the single-mutant fixation probability, and thus similar to the standard average-rate origin-fixation treatment in section IV.

#### Fixation times (Fig 6)

There are two ways in which a phenotype *p*_1_ can fix: the first scenario is that the prevalent genotype in the population is a portal genotype to *p*_1_. In this case *p*_1_ will arise repeatedly in quick succession and could fix. The second scenario is that a portal genotype exists among the wider set of less frequent genotypes in the population. Since the number of individuals with any particular genotype other than *g*_0_ is expected to be small, we do not expect *p*_1_ to arise repeatedly. Thus, the first scenario is rarer, but likely to lead to fixation when it occurs, whereas the second scenario is more common, but less likely to lead to fixation. We will thus compute and then combine the fixation time scales for both scenarios:

- **Time scale** *t*_*p*_ **for fixation from a portal genotype:** similar to origin-fixation calculations [2], this expression is a product of a ‘mutation term’ (how often a portal genotype appears) and a ‘fixation term’ (how likely *p*_1_ goes into fixation from a portal) - but unlike the NC-average equivalent, each term summarises an entire ‘burst’ of mutants, rather than a single mutation.
  – How often does the population localise on a portal genotype through neutral drift? The time until the population localises on a portal genotype for a rare phenotype 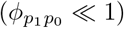 can be approximated as follows: each genotype has *L*(*K −* 1) mutational connections and one in 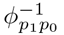 mutations from the NC give *p*_1_. Then, on average one in 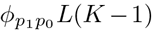 genotypes on the NC is a portal genotype to *p*_1_. Since the prevalent genotype changes every *t*_*ne*_ mutations and a portal genotype appears every 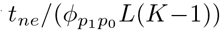 generations.
  – How likely is *p*_1_ to go into fixation from a portal? This is given by 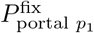 in Eq 18.
- **Time scale** *t*_*c*_ **for fixation from a wider set of genotypes:** The part of the population that forms a cloud around *g*_0_ is made up of a diverse set of genotypes, so we use an average-rate treatment, where the fixation time scale is given by Eq 13. We only need to correct for the fact that only a fraction (1 *− f*_0_) of the population is part of this cloud. This gives 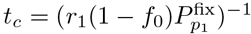

Thus, we have two different mechanisms, by which *p*_1_ could fix: either from a portal genotype on a time scale *t*_*p*_ or from a wider set of genotypes on a time scale *t*_*c*_. The time scale at which at least one of these mechanisms has taken place and *p*_1_ has gone into fixation if we assume the two mechanisms to be independent is 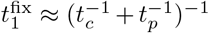(shown in the SI section 5.3). Putting this all together gives:

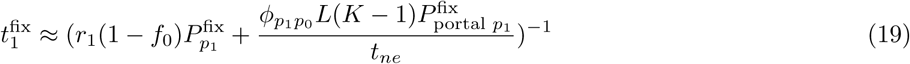

We can understand this expression if we study its limiting behaviour:

1. If *t*_*c*_ *≪ t*_*p*_, then most fixations take place from the wider cloud of genotypes and we have a fixation time of

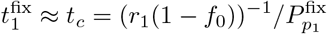

In this case, the behaviour is identical to the NC-average model, except for the correction factor (1*−f*_0_), and therefore the selective advantage of *p*_1_ has a big impact.
2. If *t*_*c*_ ≫ *t*_*p*_, most fixations take place when the prevalent genotype *g*_0_ is a portal genotype to *p*_1_. In this case, the bursty and overdispersed aspect of variation dominates the outcomes and we have

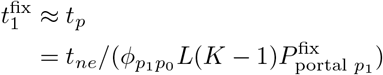

In this case, the time only depends on the selective advantage via 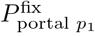, which is not only sub-linearly affected by changes in the selective advantage if the burst size is large enough (see Eq 18 and Fig 8).

#### Fixation probability of two competing mutants (Fig 6)

As before, in the NC-average model, we simply estimate the fixation times for both mutants and then calculate the probability that *p*_*r*_ fixes first as 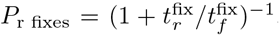. The only difference is that we use our random-map calculations (Eq 19) to compute these fixation times, rather than relying on the average-rate calculations (Eq 13).

## V. SUPPORTING CITATIONS

References [75–80] appear in the Supporting Material.

## VII. ACKNOWLEDGEMENTS

The authors acknowledge useful discussions with David McCandlish, Sam von der Dunk and Malvika Srivastava.

## VIII. AUTHOR CONTRIBUTIONS

Conceived and designed the analysis: N.S.M., S.S., C.Q.C., A.A.L.; Collected and analysed the data: N.S.M., S.S.; Wrote the paper: N.S.M., A.A.L.

## IX. DATA AVAILABILITY

The code behind this analysis can be found at https://github.com/noramartin/evolutionary_dynamics.

## Electronic supplementary material

### 1 Additional data on overdispersed variation in RNA

#### 1.1 Existence of bursts for a range of initial conditions and target phenotypes

In the main text, we only plotted the overdispersed statistics for one phenotypic transition in Fig 4. Here, we show this data for further phenotypic transitions from the same initial NC (Fig S1). We restrict ourselves to phenotypes that appear at intermediate rates 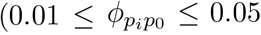, excluding the unfolded structure) since these appear at sufficiently high rates to collect sufficient data for a bar chart, but at sufficiently low rate that they are not produced continuously. We find that the appearance of new phenotypes *p*_*i*_ is overdispersed for all these choices of *p*_*i*_.

**Figure S1:**
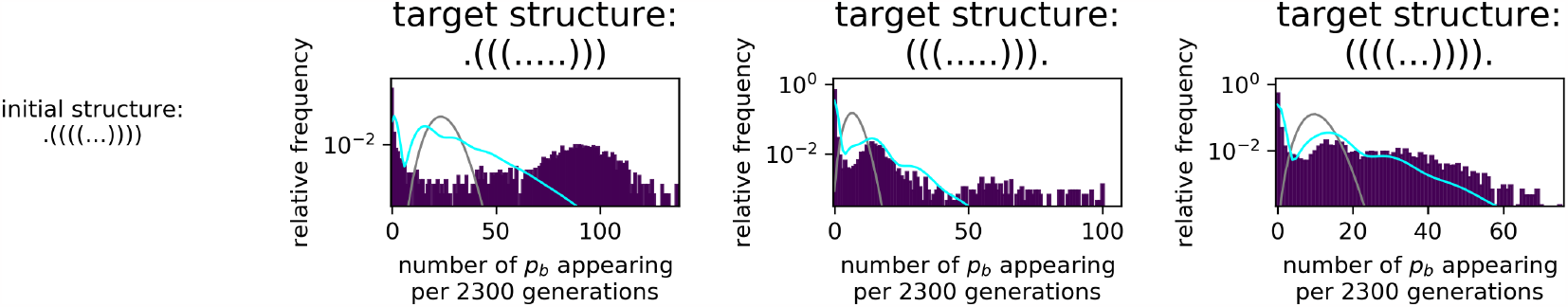
Strong deviations from Poisson statistics for the appearance of phenotype p_b_ in a population neutrally evolving with stabilising selection for p_g_. As in Fig 4 in the main text, the appearance of novel phenotypes is quantified by splitting the simulation into time intervals of Δt = 2300 generations and recording how often a given new phenotype p_b_ appears in each Δt. This data is for the RNA GP map only. The number of appearances per interval is highly overdispersed compared to a Poissonian distribution with the same mean (grey line), which would be observed in the average-rate model. The overdispersion is also higher than analytically predicted for the random map (cyan line), which is not surprising given that the RNA GP map includes non-neutral correlations that do not exist in the random map. Parameters: population size N = 1000, mutation rate u = 2 × 10^−5^, total time 10^7^ generations. The initial NC is the one shown in Fig 1 in the main text, and the target phenotypes p_b_ are given above each plot in dot-bracket notation.

#### 1.2 Dependence of burstiness on population parameters for a range of initial conditions

**Figure S2:**
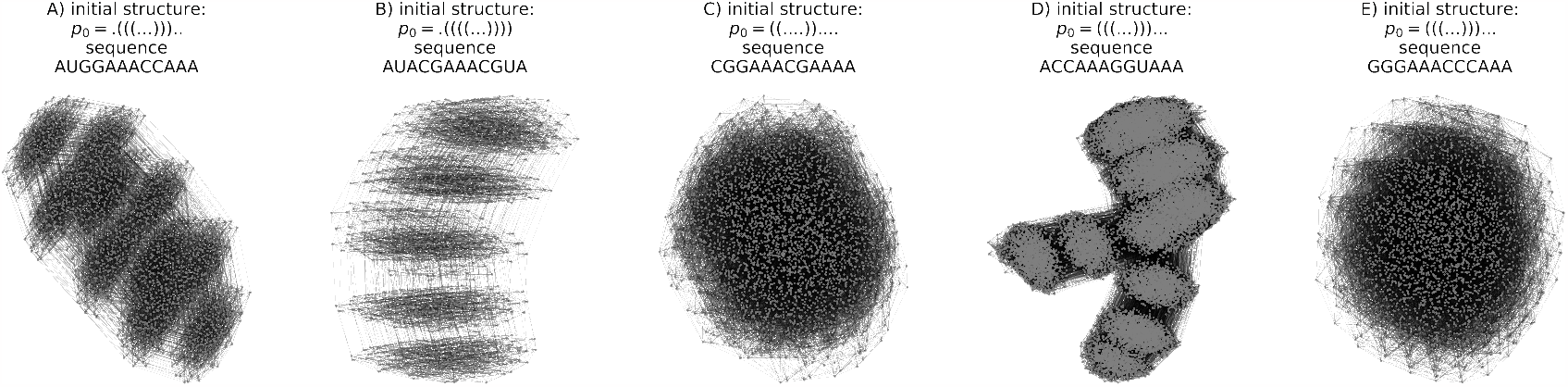
The NCs used as starting points in the simulations of evolving populations: these five NCs were randomly chosen from all large NCs (with > 10^3^ genotypes). Each NC is one connected network within the neutral set of a given phenotype, and is uniquely defined once we know one sequence in the NC (then any other sequence that can be reached through a series of neutral mutations from that sequence is part of the NC). Thus, the title of each network includes both the RNA structure associated with the neutral set in dot-bracket notation, and one sequence to identify the NC. The NC in Figs 1-6 of the main text is the second one. The graphs are drawn using the force-layout algorithm from NetworkX [1].

In the main text, we quantified the burstiness of the appearance of a new phenotype *p*_*i*_ by the coefficient of variation of the time intervals between two appearances: if *p*_*i*_ phenotypes appear in a constant-rate Poisson process, this coefficient would be one, but if *p*_*i*_ phenotypes appear in bursts, it is greater than one. We plotted this coefficient of variation a range of population parameters in the main text, but limited ourselves to a single initial NC and a single target phenotype. Here, we repeat the same analysis for populations starting at all five NCs from Fig S2. These five initial NCs, which include the NC from the main text, are randomly chosen from all large NCs (with *>* 10^3^ genotypes). We only consider large NCs for the following reasons:

- While small and large NCs can have phenotypic transitions with high *ϕ*_*pq*_, only large NCs have phenotypic transitions with small *ϕ*_*pq*_ since low-frequency transitions can only exist if they occur a small number of times in a large NC. Thus, large NCs are needed to explore phenotypic transitions with a range of *ϕ*_*pq*_ values.
- The randomisation employed in creating the different models essentially shuffles the mutational neighbourhood of a NC and this is more effective in large NCs.
- Because phenotypes with large neutral sets are more likely to appear in nature [2–4], large neutral sets and therefore large NCs are likely to play an important role in evolution.
- Neutral set sizes and NCs for longer sequence lengths are likely to be even larger, so it is important to understand the dynamics on large NCs.

For each of these initial NCs, we studied the overdispersion for a variety of target phenotypes for each NC: this data is shown in Figs S3 - S7, one figure for each initial NC. For each of the five starting NCs, the set of target phenotypes was chosen according to the following criteria: we need phenotypes that are frequent enough that they appear sufficiently often in the simulation to estimate their statistics. However, we also need them to be rare enough that they are unlikely to appear in every generation since our analysis relies on the mean and standard deviation of the time intervals between two appearances, and this analysis assumes continuous time intervals, which is not a good assumption if these time intervals are frequently one or even zero generations (especially since two concurrent appearances in the same generation would be counted as one appearance in this analysis). Therefore, data was collected for all target phenotypes with intermediate frequency (0.01 *≤ ϕp*_*i*_*p*_0_ *≤* 0.05). Out of this set, we applied additional criteria before including a given target phenotype in the plot: we checked that the phenotype appears at least 500 times in every simulation, and less frequently than every 5 generations, to ensure sufficient data and to avoid problems with discrete generation times.

The results in Figs S3 - S6 are in agreement with those shown in the main text: the overdispersion tends to be weakest on the random map and strongest on the RNA GP map due to its highly inhomogeneous structure. Furthermore, the overdispersion tends to decrease with increasing genetic drift (small population size *N*) or increasing genetic diversity (large *uLN*), consistent with the scaling arguments in the main text and ref [5]. Note that it is difficult to capture a complex phenomenon like overdispersion in a simple parameter: there are limitations in quantifying burstiness of finite data sets using simply metrics based on the coefficient of variation (see for example [6]); and thus small quantitative differences in the coefficient of variation may not be significant, and reliable conclusions can only be reached by also investigating the full distributions, as we did in Fig 4 in the main text and Fig 1.1 in this document.

**Figure S3:**
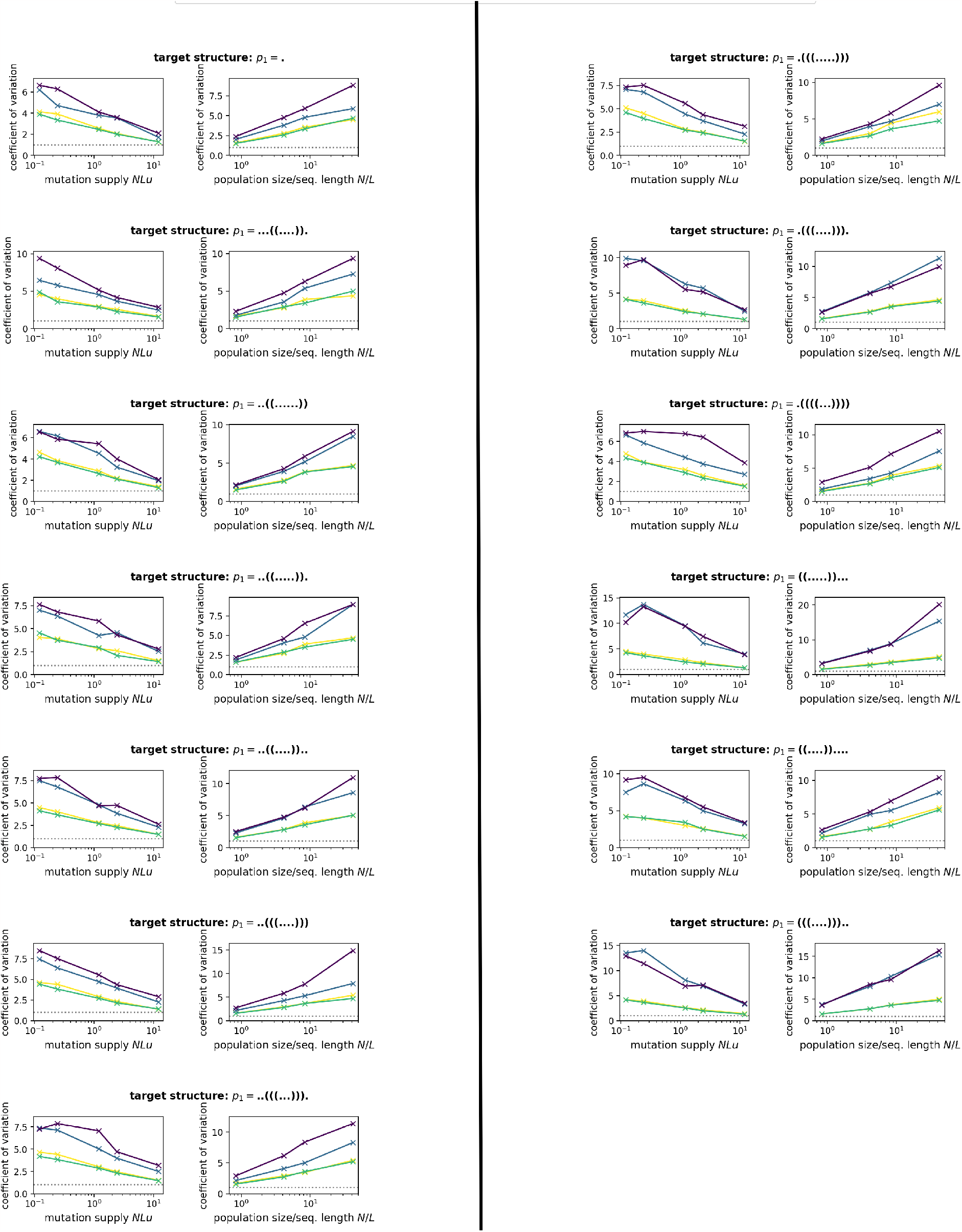
How does the amount of overdispersion, quantified by the coefficient of variation, depend on the population parameters?: this figure shows the same analysis as Fig 5 in the main text: as a measure of the burstiness, the coefficient of variation of the time distribution between two p_i_ appearances is computed for populations with a range of population parameters. Here, the initial NC and phenotype is the one in Fig S2A. The target phenotype p_b_ is a different one in each row and indicated in the row title. For each target phenotypes, there are two analyses: (first column) a range of mutation rates u for a fixed population size of N = 200 and (second column) a range of mutation rates N for a fixed mutation rate of u = 5 × 10^−5^.

**Figure S4:**
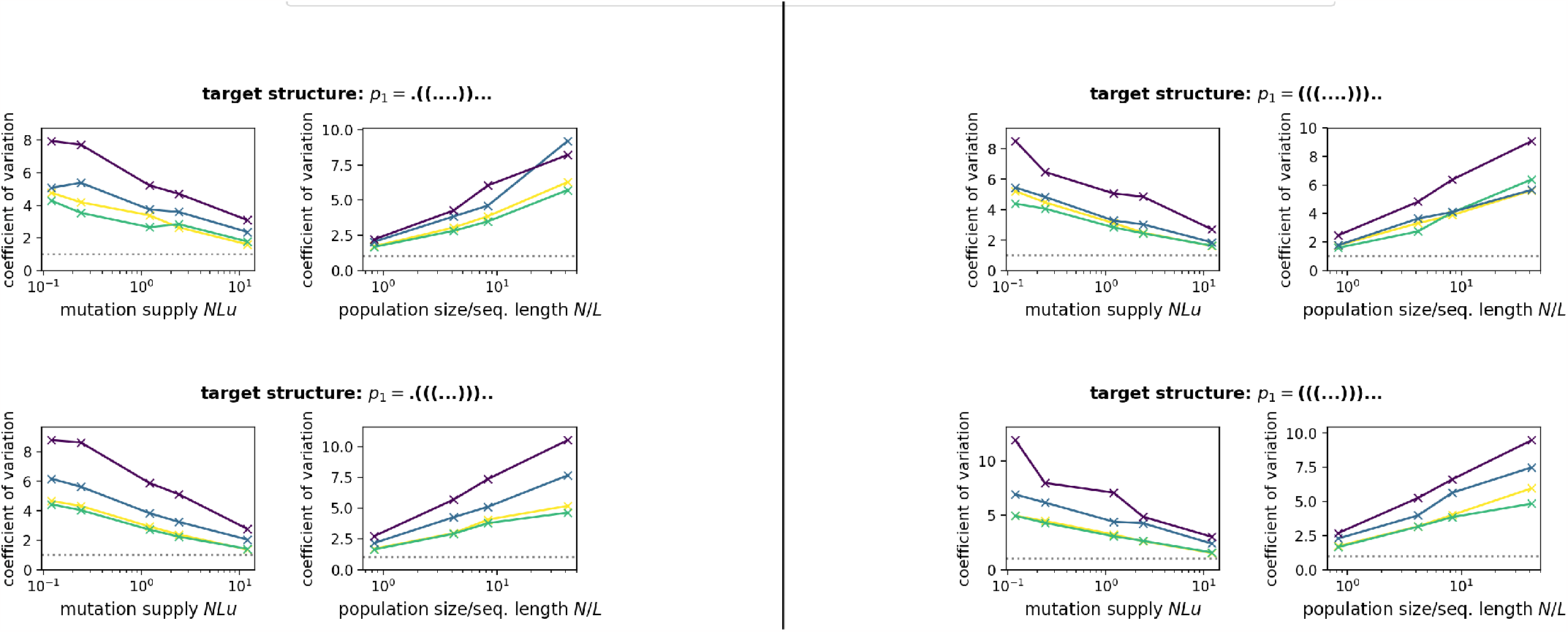
Same as Fig S3, but here the initial NC and phenotype is the one in Fig S2B.

**Figure S5:**
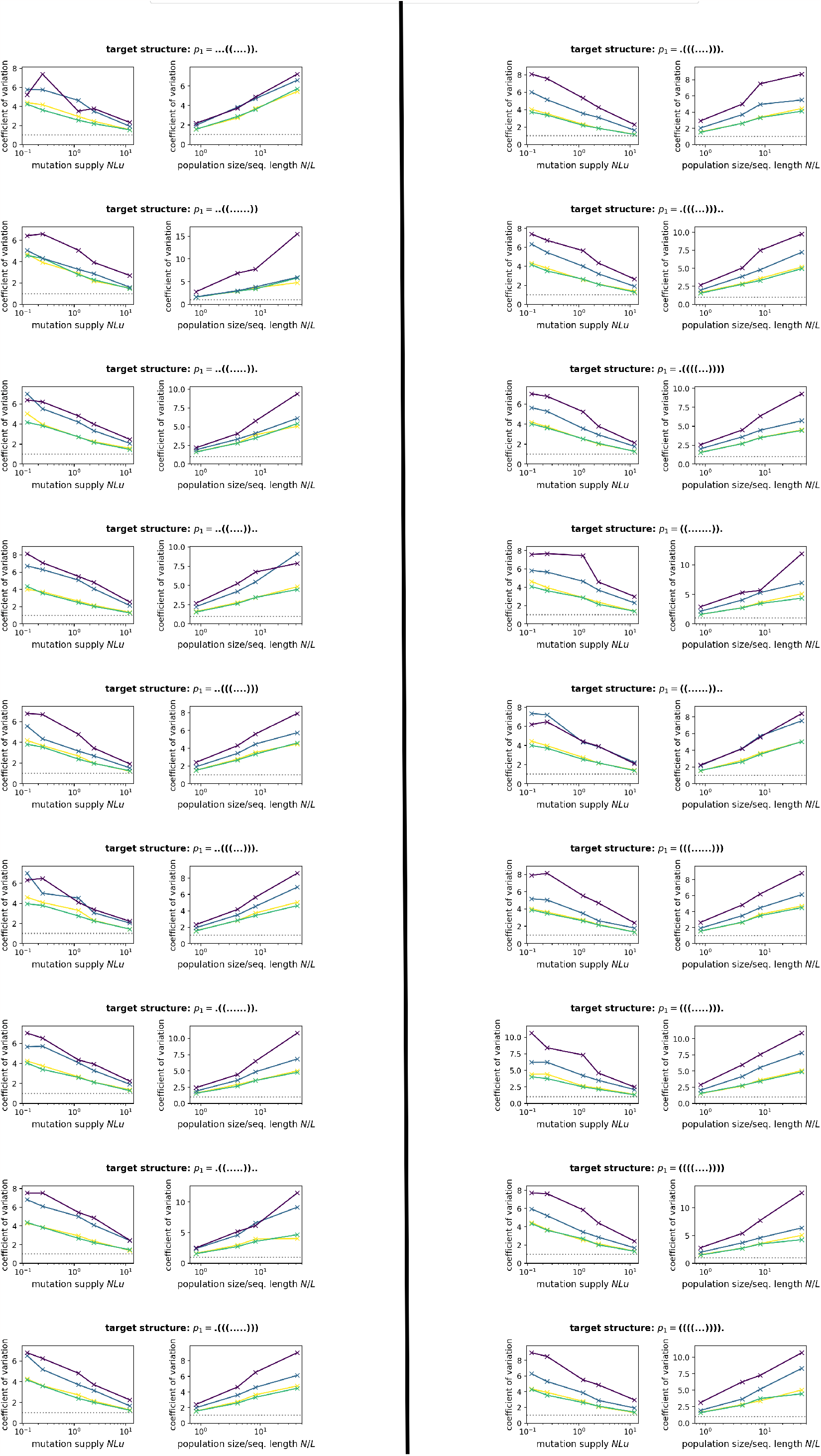
Same as Fig S3, but here the initial NC and phenotype is the one in Fig S2C.

**Figure S6:**
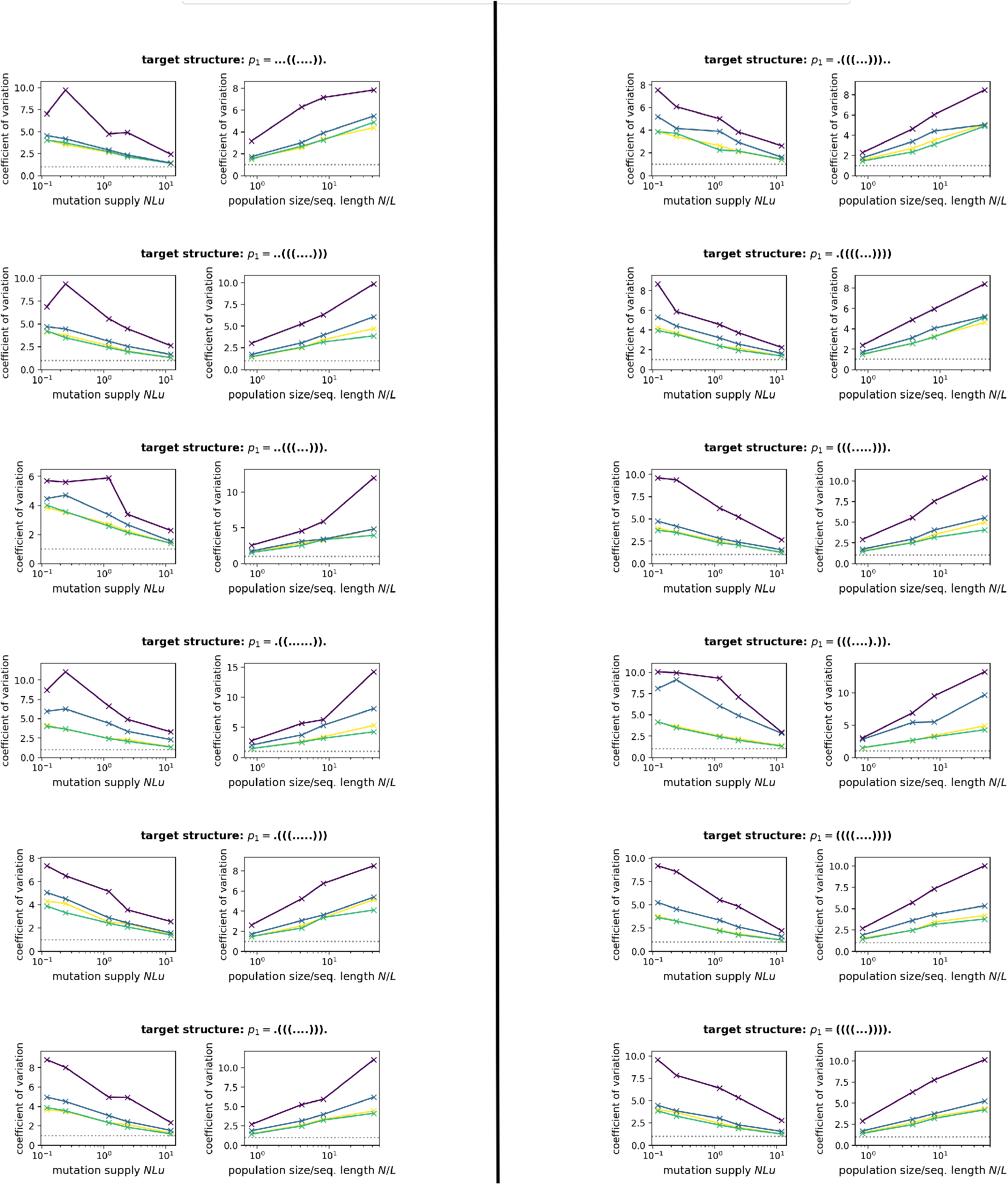
Same as Fig S3, but here the initial NC and phenotype is the one in Fig S2D.

**Figure S7:**
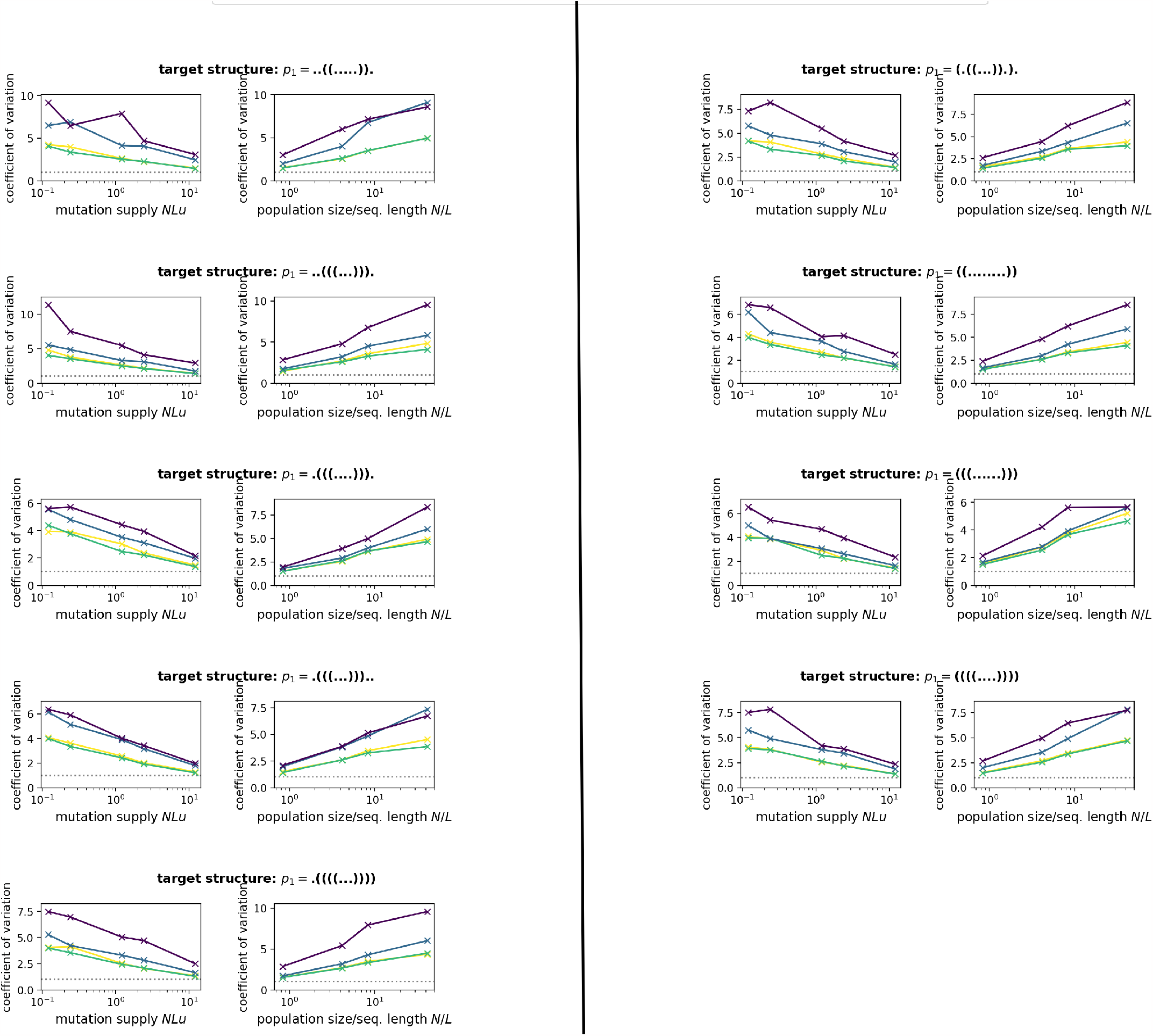
Same as Fig S3, but here the initial NC and phenotype is the one in Fig S2E.

#### 1.3 Longer sequence length of *L* = 30

**Figure S8:**
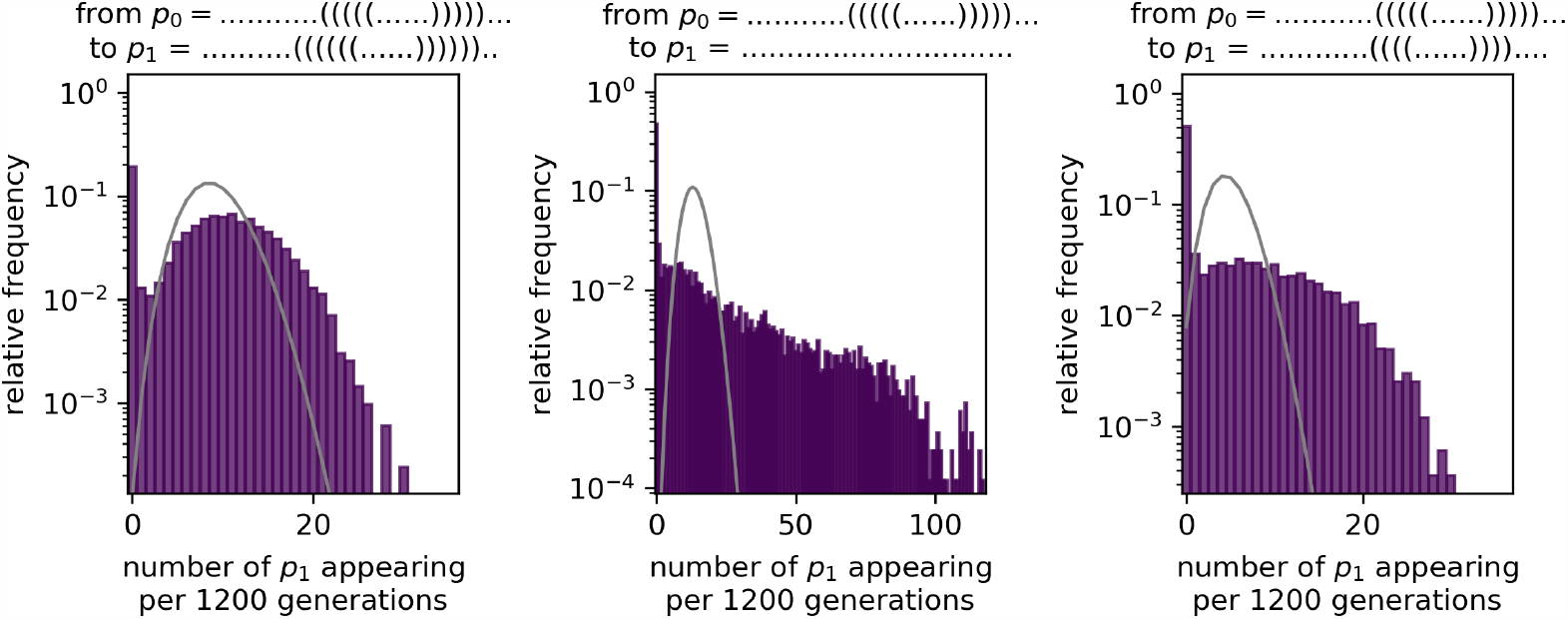
Overdispersion in an evolutionary process with RNA structures of thirty nucleotides in length: as in Fig 4 in the main text, we simulated a population under strong stabilising selection, and counted how many times a new phenotype p_b_ is introduced in a time interval of Δt = 1600 generation. We find that p_b_ appears in a bursty manner compared to a Poissonian distribution with the same mean (grey line), which would be expected in a constant-rate process. Parameters: population size N = 1000, mutation rate u = 1×10^−5^, total time 10^7^ generations. In each subplot, the initial and final structure are given in the plot titles - the initial structure p_0_ is the same throughout this plot since all data is derived from a single evolutionary simulation, where selection stabilises this specific structure. For final structure p_1_, we restricted ourselves to structures that are expected to appear with intermediate frequencies of 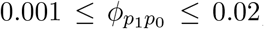, since our simulations are not long enough to collect sufficient data for low-frequency structures (we only plot data for structures that appear at least 10^3^ times and on average more than twice in each Δt). To apply this criterion, we approximate the 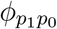values using the procedure from our previous work [7], which uses the site-scanning sampling method [8].

**Figure S9:**
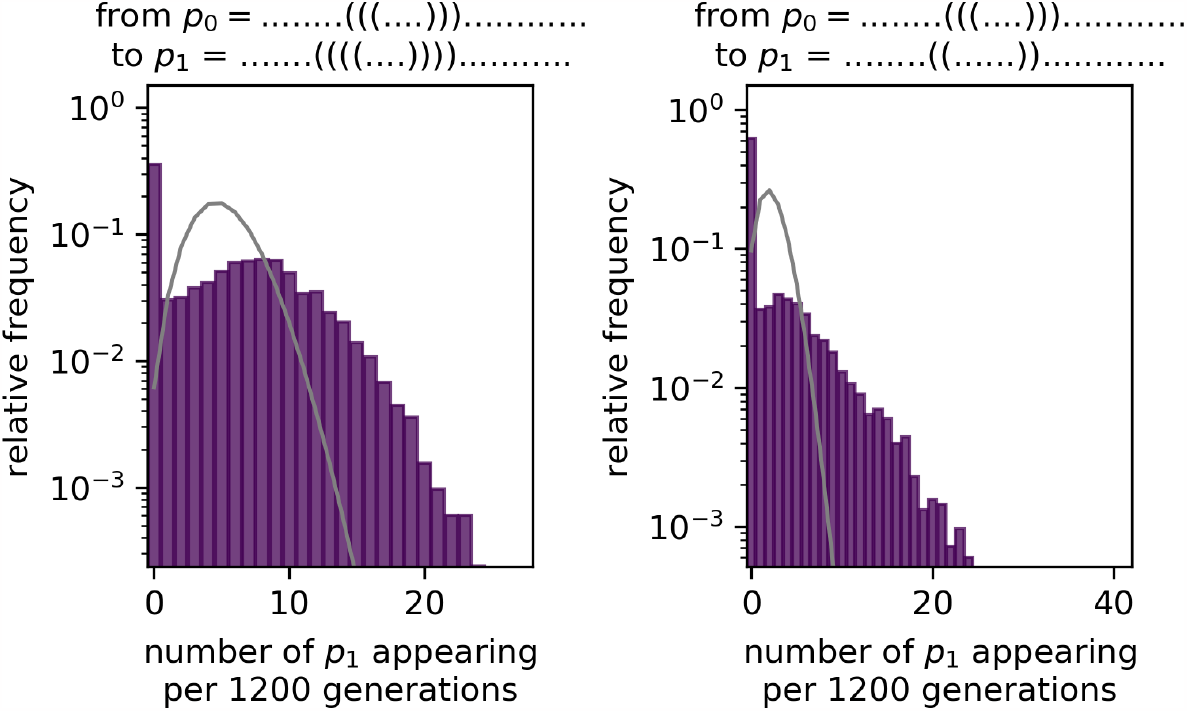
Overdispersion in an evolutionary process with RNA structures of length L = 30: same as Fig S8, but for a different initial structure p_0_.

Throughout the main text, we worked with RNA sequences of length *L* = 12 and their secondary structures. This allowed us to enumerate and fold all possible sequences of that length, visualise the entire neutral spaces and create null models with the same statistics. Such a full analysis is not feasible for much longer sequence lengths, since the number of sequences that would have to be folded to gain a full overview of the GP map grows exponentially as 4^*L*^. However, sophisticated sampling methods have shown that NCs still have a pronounced community structure for sequences up to *L* = 45 [9], indicating that NCs remain inhomogeneous, and non-neutral correlations have been shown for sequences up to *L* = 20 [10]. Even without this imhomogeneous structure, on the random map, we expect overdispersion under the conditions outlined in the main text.

**Figure S10:**
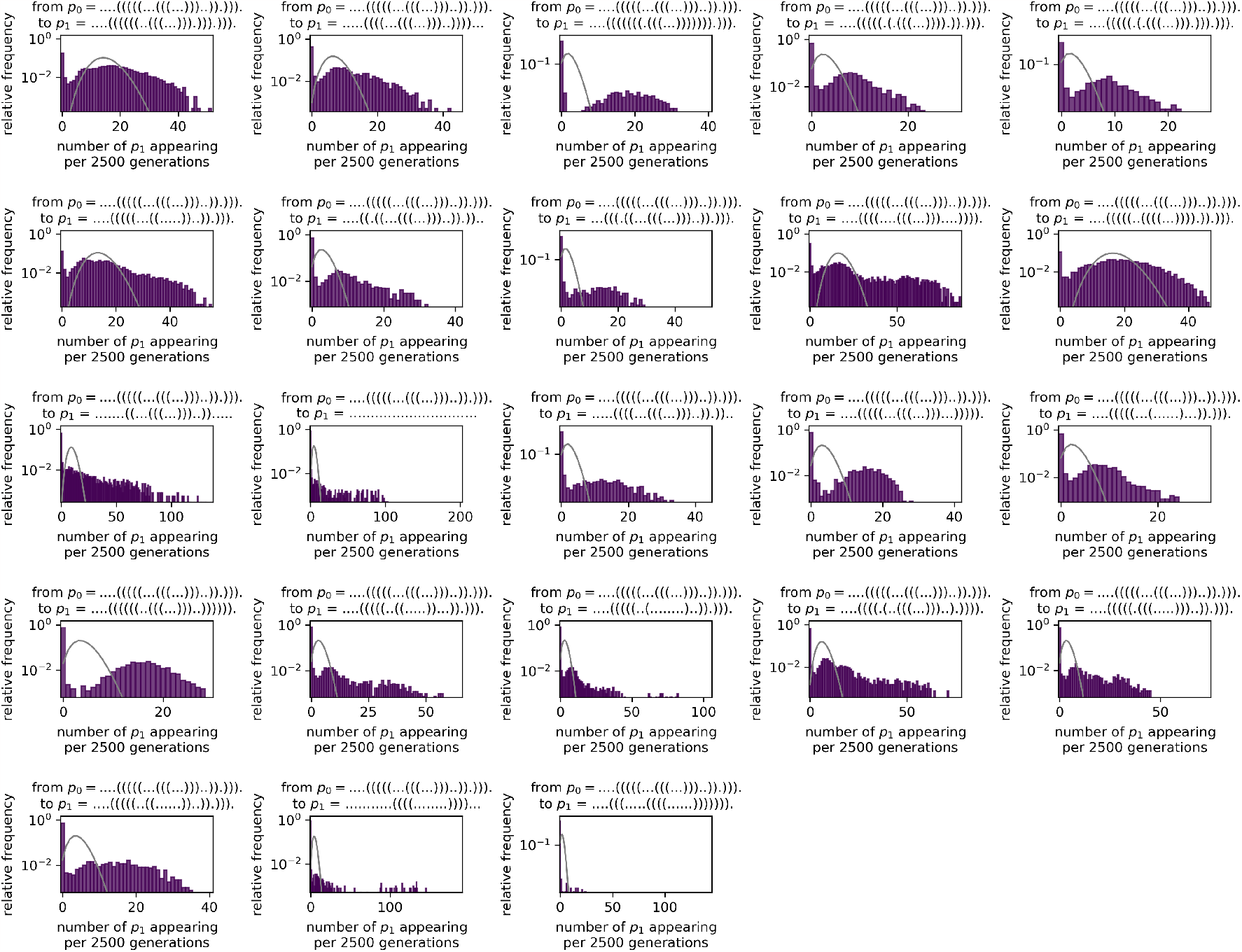
Overdispersion in an evolutionary process with RNA structures of length L = 30: same as Fig S8, but for a different initial structure p_0_.

**Figure S11:**
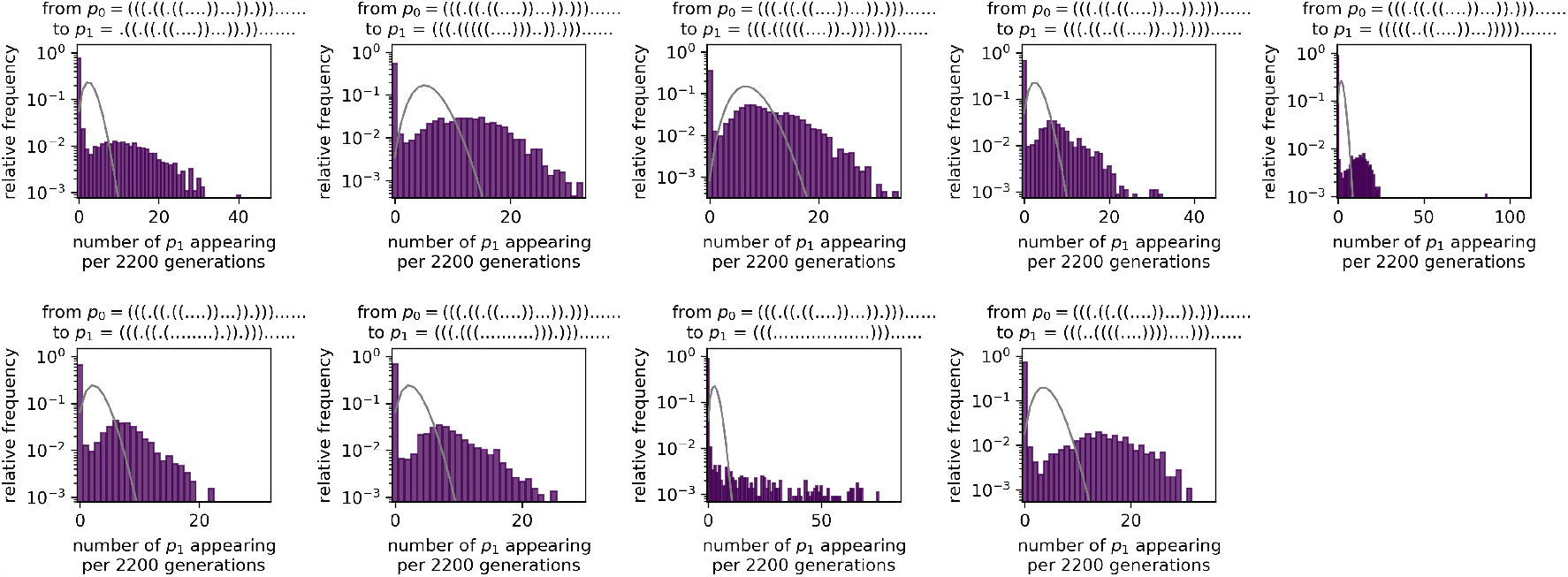
Overdispersion in an evolutionary process with RNA structures of length L = 30: same as Fig S8, but for a different initial structure p_0_.

**Figure S12:**
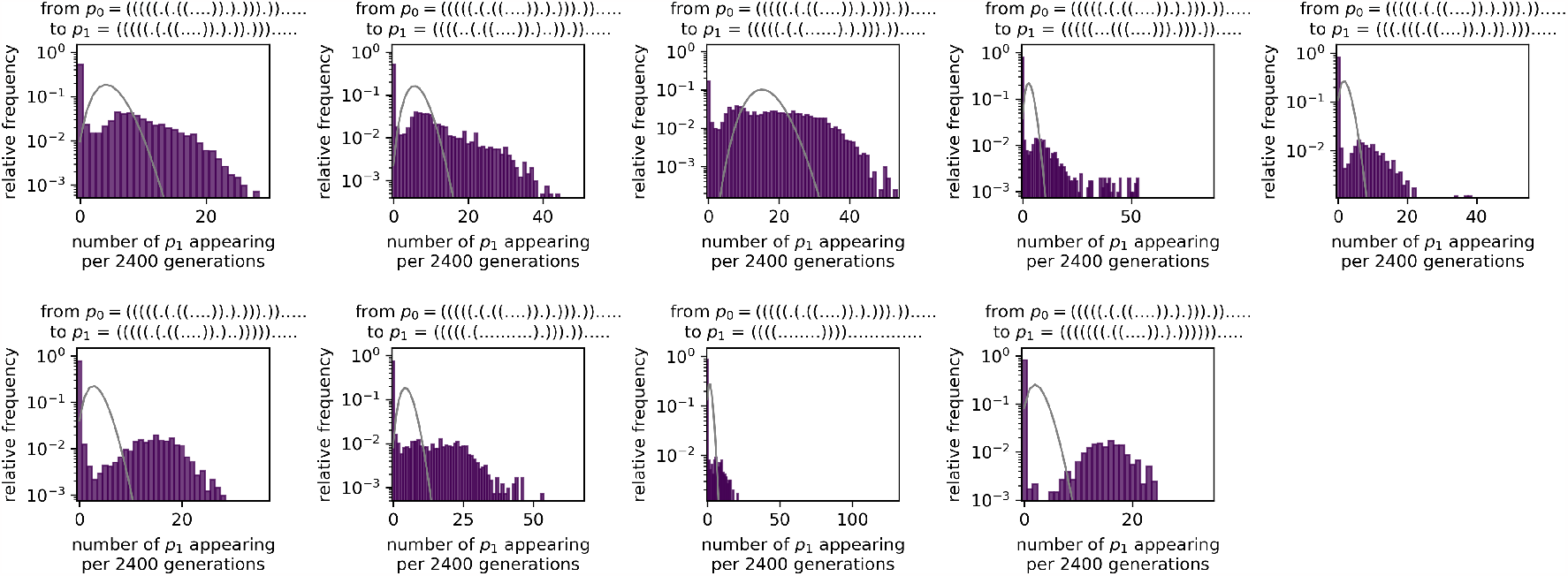
Overdispersion in an evolutionary process with RNA structures of length L = 30: same as Fig S8, but for a different initial structure p_0_.

Here, we can simulate evolutionary processes for sequences of length *L* = 30, which is too long for an exhaustive analysis since the number of sequences of that length is 4^30^ *≈* 10^18^. Figs S8 - S12 shows that the key result from the main text remains unchanged: new phenotypes *p*_*i*_ appear in overdispersed bursts. These Figures show data for initial structures with between one and four stacks - since the number of stacks in a structure is closely linked to its neutral set size [4] (and thus mutational robustness [10] and folding stability [11]), including a range of number of stacks ensures a diverse set of structures (as in our previous work [7]).

#### 1.4 Theoretical predictions of fixation times for *L* = 50 and *L* = 500

To further investigate, in how far our results generalise to longer sequences, we apply our analytic approximations for fixation times to RNA sequences of length *L* = 50 and *L* = 500 (Fig S13). As before, we find that fixation times can differ by many orders of magnitude between the average-rate prediction and the prediction for the random GP map.

**Figure S13:**
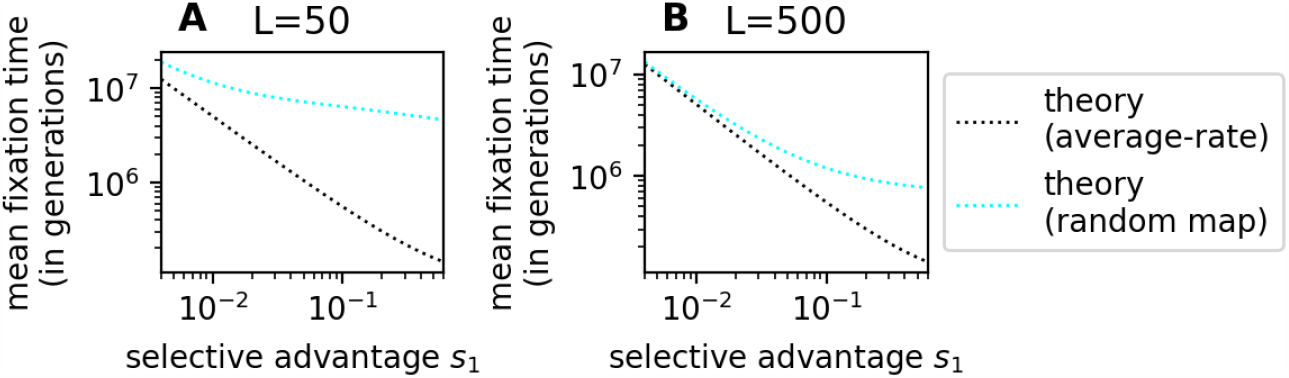
The selective advantage s_1_ of an adaptive phenotype has a weaker influence on its fixation time in the overdispersed case: these plots shows analytic predictions for RNA sequences of length (A) L = 50 and (B) L = 500, modelling the following scenario: the population starts with an initial phenotype p_0_, a single phenotype p_1_ has a selective advantage of s_1_ over p_0_ and all other phenotypic changes are deleterious with zero fitness. We predict analytically, how many generations it takes on average until p_1_ fixes for two cases: first, we assume that p_1_ appears at constant rate (grey line), and secondly, we repeat the calculation for the corresponding random GP map (teal line), where there are several discrete genotypes in each neutral space and each genotype’s mutational neighbourhood is drawn from a fixed distribution. In all cases, p_1_ fixes more rapidly if its selective advantage is higher, but this decrease is steeper for the constant-rate model than in the random map calculation, which has overdispersed dynamics. Parameters: population size N = 10^3^, mutation rate u = 10^−4^/L (to keep the mutation supply constant across the figure), average rate of mutating to 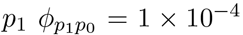, robustness of the initial phenotype 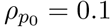. These parameters lead to burst sizes of M = N/((K − 2)ρL) of 66.7 for L = 50, and N = 6.67 for L = 500. As expected, significant deviations from the average-rate model emerge for selection coefficients larger than about 1/M. The lowest selective advantage considered is 4/N since we are not interested in effectively neutral mutations with s ⪅ 1/N.

### 2 Overdispersed variation in other GP maps

#### 2.1 Overdispersed variation in the HP model for protein tertiary structure

**Figure S14:**
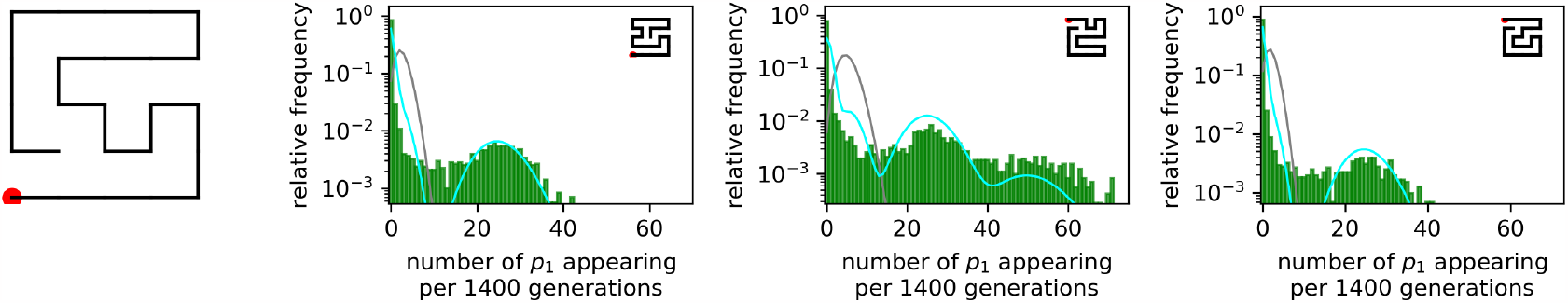
Overdispersion in an evolutionary process for HP model proteins: as in Fig 4 in the main text, we simulated a population under strong stabilising selection, and counted how many times a new phenotype p_b_ is introduced in a time interval of Δt = 1400 generation. Our results show that p_b_ introductions are highly overdispersed compared to a Poissonian distribution with the same mean (grey line), which would be observed in a constant-rate process. The analytic prediction for the random map is shown in cyan. Parameters: population size N = 1000, mutation rate u = 2 × 10^−5^, total time 10^7^ generations. The initial structure p_0_ (shown in the first subplot) is the same throughout this plot since all data is derived from a single evolutionary simulation with stabilising selection for this initial structure. The target structure for each subplot is shown in the top-right corner. We restricted ourselves to target structures that are expected to appear with intermediate frequencies 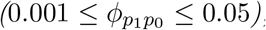, since our simulations are not long enough to collect sufficient data for low-frequency structures (we only plot data for structures that appear at least 500 times and on average at least twice in each Δt).

**Figure S15:**
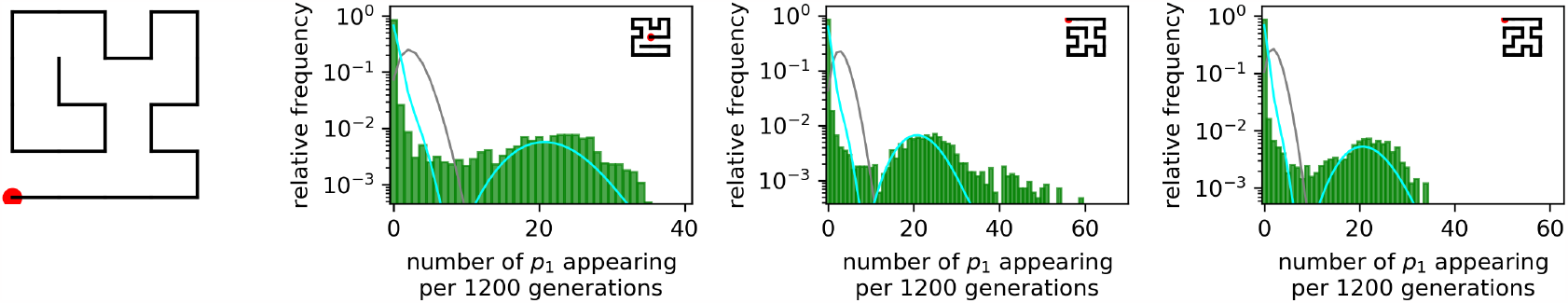
Overdispersion in an evolutionary process for HP model proteins: same as Fig S14, but for a different initial structure (shown in the first subplot).

**Figure S16:**
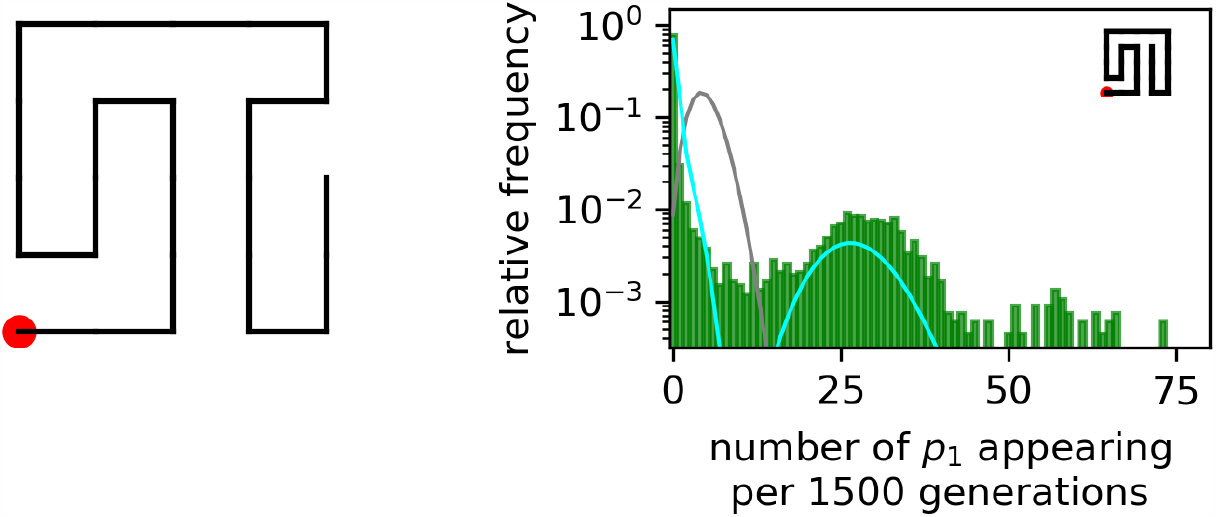
Overdispersion in an evolutionary process for HP model proteins: same as Fig S14, but for a different initial structure (shown in the first subplot).

**Figure S17:**
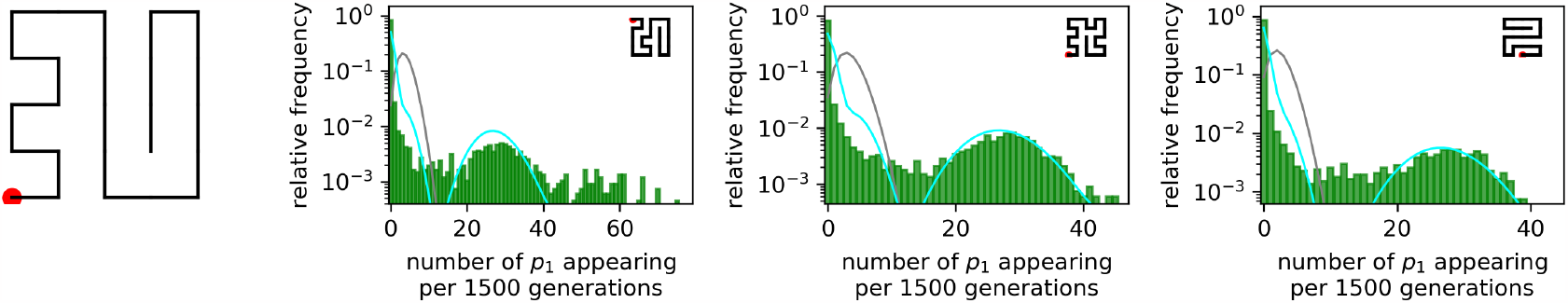
Overdispersion in an evolutionary process for HP model proteins: same as Fig S14, but for a different initial structure (shown in the first subplot).

Because proteins and their folded structures play a central role in molecular biology, we also test our hypothesis on a simple model of protein tertiary structure, the well-studied [10, 12–14] HP model. This model simulates molecular chains on a lattice. The chains consist of hydrophobic and polar residues, fold into a structure that maximises the number of contacts between hydrophobic residues and thus simulate one important aspect of protein folding. Here, we use the Python implementation and data for compact 5 *×* 5 HP lattice proteins from our previous work [7]. Sequences with several degenerate mfe structures are treated as non-folding and not considered further.

As before for the RNA map, we find that phenotypic variation in the HP protein model is overdispersed (see Figs S14 - S17). However, unlike for RNA, this overdispersion is not much higher than that expected for a random map without genetic correlations. This is consistent with previous observations that non-neutral correlations, which would amplify the bursts compared to the random map, are weaker in the HP protein model than in the RNA GP map [10]. Whether this is an artefact of the coarse-grained nature of the HP model and its energetics, or whether non-neutral correlations are truly weaker in proteins than in RNA, is a question for future research.

#### 2.2 Overdispersed variation in Richard Dawkins’ biomorphs, a toy model of development

**Figure S18:**
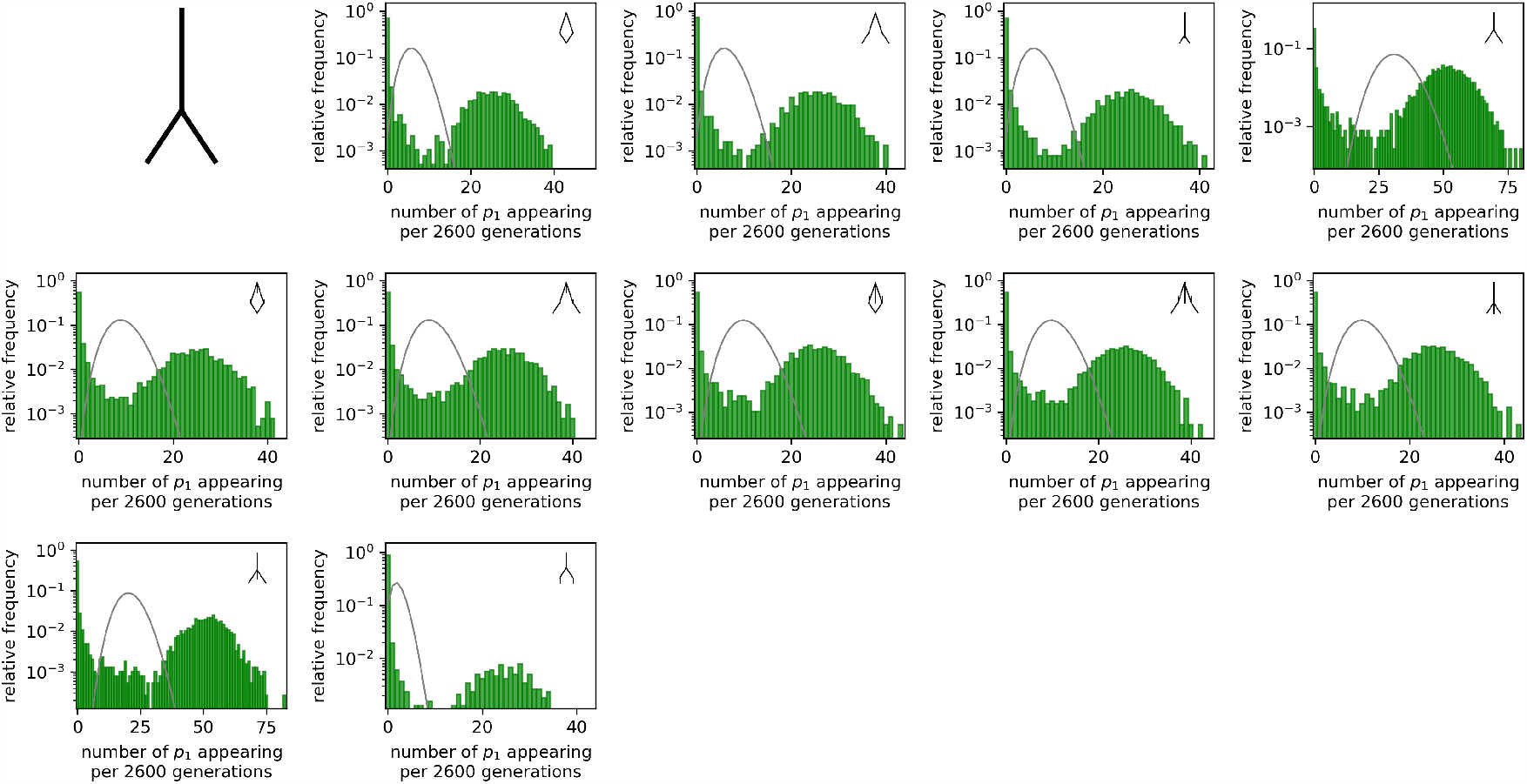
Overdispersion in an evolutionary process for Richard Dawkins’ [15] biomorphs: as in Fig 4 in the main text, we simulated a population under strong stabilising selection, and counted how many times a new phenotype p_b_ is introduced in a time interval of Δt = 2600 generation. Our results show that p_b_ introductions are highly overdispersed compared to a Poissonian distribution with the same mean (grey line), which would be observed in a constant-rate process. Parameters: population size N = 1000, mutation rate u = 2×10^−5^, total time 10^7^ generations. The initial phenotype p_0_ (shown in the first subplot) is the same throughout this plot since all data is derived from a single evolutionary simulation, where only this initial phenotype is viable. The target phenotype for each subplot is shown in the top-right corner. We restricted ourselves to target structures that are expected to appear with intermediate frequencies 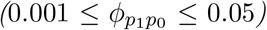, since our simulations are not long enough to collect sufficient data for low-frequency structures (we only plot data for structures that appear at least 500 times and on average at least twice in each Δt).

As one second test, we work with a GP map that is based on completely different principles than the molecular RNA and HP models explored so far: Richard Dawkins’ biomorphs. Biomorphs are a simple toy model of development and are based on recursive branching rules, which imitate growth processes in nature [15]. Biomorphs have a genotype, which is made up of nine integer values, and map this to a phenotype, a 2D drawing. This 2D drawing can be converted into a discrete, clearly defined phenotype, using methods in ref [16].

Here, we repeat our analysis from the main text for the biomorphs GP map to investigate, whether we observe bursts in evolutionary processes in the biomorphs system. Fig S18 shows that we do indeed find such bursts. Thus, bursts are not only found in evolutionary processes on molecular GP maps, but also in the biomorphs system as one example of a developmental model based on recursive growth processes.

### 3 Effect of quasi-simultaneous *p*_1_ mutants during a burst

In all of our calculations and arguments in this paper, we have used the single-mutant fixation probability for each new phenotypic introduction. However, in the bursty case, *p*_1_ mutants can arise at high rates during a burst, which greatly increases the likelihood that two *p*_1_ mutants arise in quick succession or even in the same generation. To understand if this has any effect on fixation processes, we need to compare the probability of fixation for two limiting cases: (I) *n* phenotypes *p*_1_ undergo independent fixation or extinction processes (i.e. one *p*_1_ mutant either fixes or is lost before the next one appears) or (II) *n* phenotypes *p*_1_ appear in a single generation.

#### 3.1 Analytic treatment using Kimura’s fixation probability

First, let us investigate the difference between these two cases using the classic expression for the probability of fixation of *n p*_1_ phenotypes with a selective advantage of *s*_1_ in a population of *N* haploid organisms [17]:

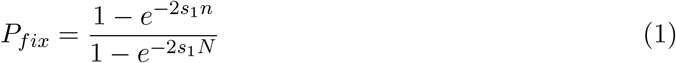

Let us denote the denominator of Equation 1 as:

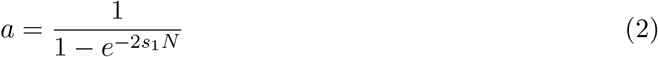

This constant *a* is approximately ∼ 1 in the limit of strong selection (*Ns*_1_ *≫* 1) and *≪* 1 for weak selection (*Ns*_1_ *≪* 1).

Then we can write the probability of fixation for case (I), where *n* phenotypes *p*_1_ undergo independent fixation or extinction processes, as:

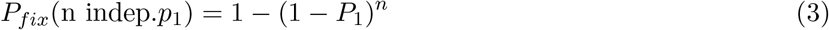

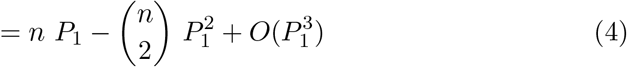

Here *P*_1_ is the probability of fixation for a single mutant.

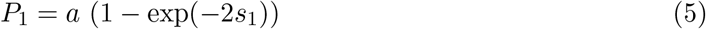

On the other extreme, the probability of fixation for case (II), where *n* mutants of type *p*_1_ appear in the population simultaneously, is:

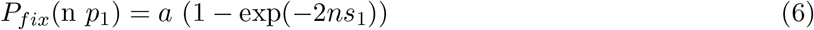

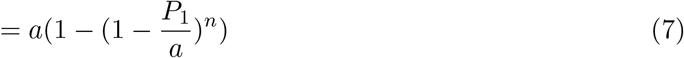

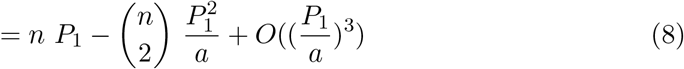

In the limit of strong selection, we have *a* ∼ 1, and so Equations 7 and 3 are identical. Therefore, the probability of fixation per mutant is the same, regardless of whether fixation processes are independent or simultaneous.

However, in the limit of weak selection, *a ≫* 1, Equations 3 and 7 are different. Their Taylor expansions, Equations 4 and 8, indicate that the fixation probability per mutant is higher when mutants arrive in groups. However, this difference is likely to be small since it scales with 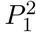, where *P*_1_ is the probability of fixation for a single mutant, which is small in the limit of weak selection.

#### 3.2 Simulations with concurrent or individual mutants

To test our analytic arguments, we simulate the fixation probabilities computationally for the two limiting cases: (I) *n* phenotypes *p*_1_ undergo independent fixation or extinction processes (i.e. one *p*_1_ mutant either fixes or is lost before the next one appears) or (II) *n* phenotypes appear in a single generation. We implement these scenarios as Wright-Fisher models without random mutations, such that all *p*_1_ mutants are introduced in a controlled fashion, either all at once in the first generation or one by one as soon as the last *p*_1_ mutant has either been lost or fixed. We repeat this setup for a range of selective advantages *s*_1_ and population sizes *N* and show the data in Fig S19. The results are consistent with our theoretical considerations: the fixation probability of *p*_1_ does not depend strongly on whether all *p*_1_ mutants are introduced at once in the first generation, or one-by-one once the last fixation process has completed. Only in the very weak-selection, quasi-neutral mutation regime (*Ns*_1_ *<* 1), can we observe a slight difference between the two scenarios: as predicted in our analytic calculations, in the quasi-neutral mutation regime, the fixation probability is higher if the mutants are introduced all at once. We can neglect this effect in the calculations in this paper since we are specifically interested in adaptation, not quasi-neutral evolution.

**Figure S19:**
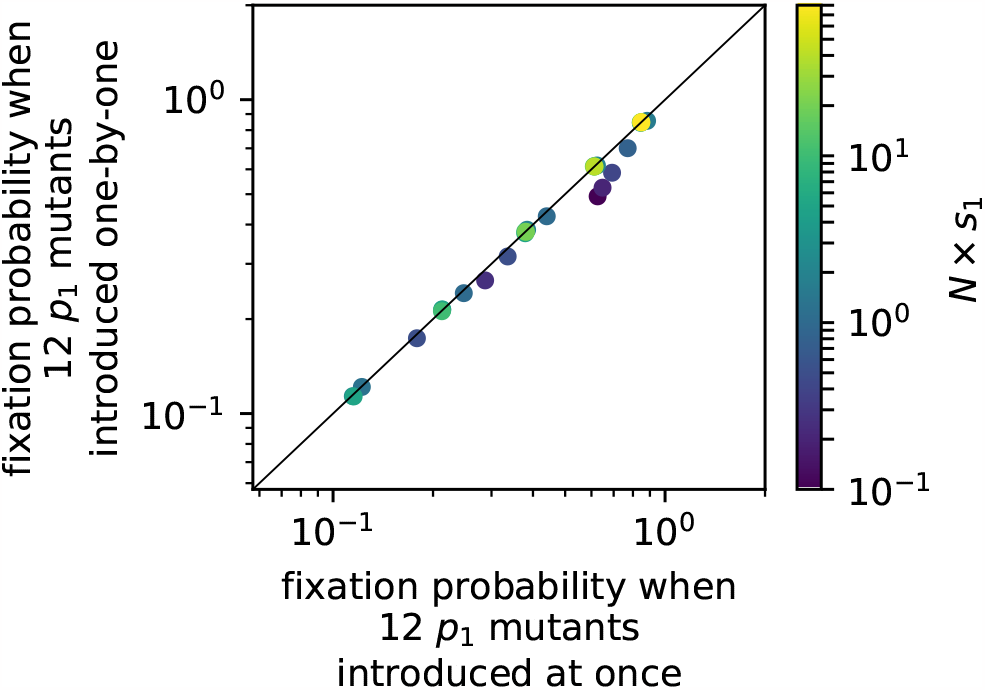
Does it make a difference whether p_1_ mutants with a selective advantage of s_1_ are introduced all at once in a single generation, or independently (i.e. after the last mutant has gone into fixation or to extinction): we consider a range of population sizes N (N = 20, N = 50, N = 100, N = 250, N = 500, N = 1000) and selective advantages s_1_ (s_1_ = 0.005, s_1_ = 0.01, s_1_ = 0.02, s_1_ = 0.04, s_1_ = 0.08) and run two sets of simulations with a Wright-Fisher model: (1) 12 phenotypes p_1_ undergo independent fixation or extinction processes (i.e. one p_1_ mutant either fixes or is lost before the next one appears) or(2) 12 phenotypes appear in a single generation. We do not introduce any further mutations and simply run the simulation until the population only consists of p_0_ or p_1_. We then compute the probability of an all-p_1_ outcome based on 10^5^ such runs. We find, consistent with our analytic considerations (eqs 1-8), that the fixation probability is the same in both cases, except in the quasi - neutral limit Ns_1_ ⪅ 1, where p_1_ is slightly more likely to fix if all p_1_ mutants appear at once.

### 4 Amount of polymorphism - theory versus simulation

Our theory for the overdispersed variation on the random GP map requires estimates of the amount of genetic diversity in an evolving population close to the monomorphic limit. In the main text, we derived the following approximation for the fraction of individuals *f*_0_ that share the prevalent genotype:

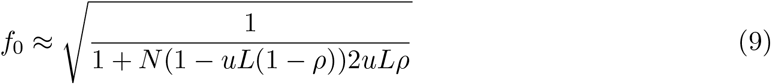

**Figure S20:**
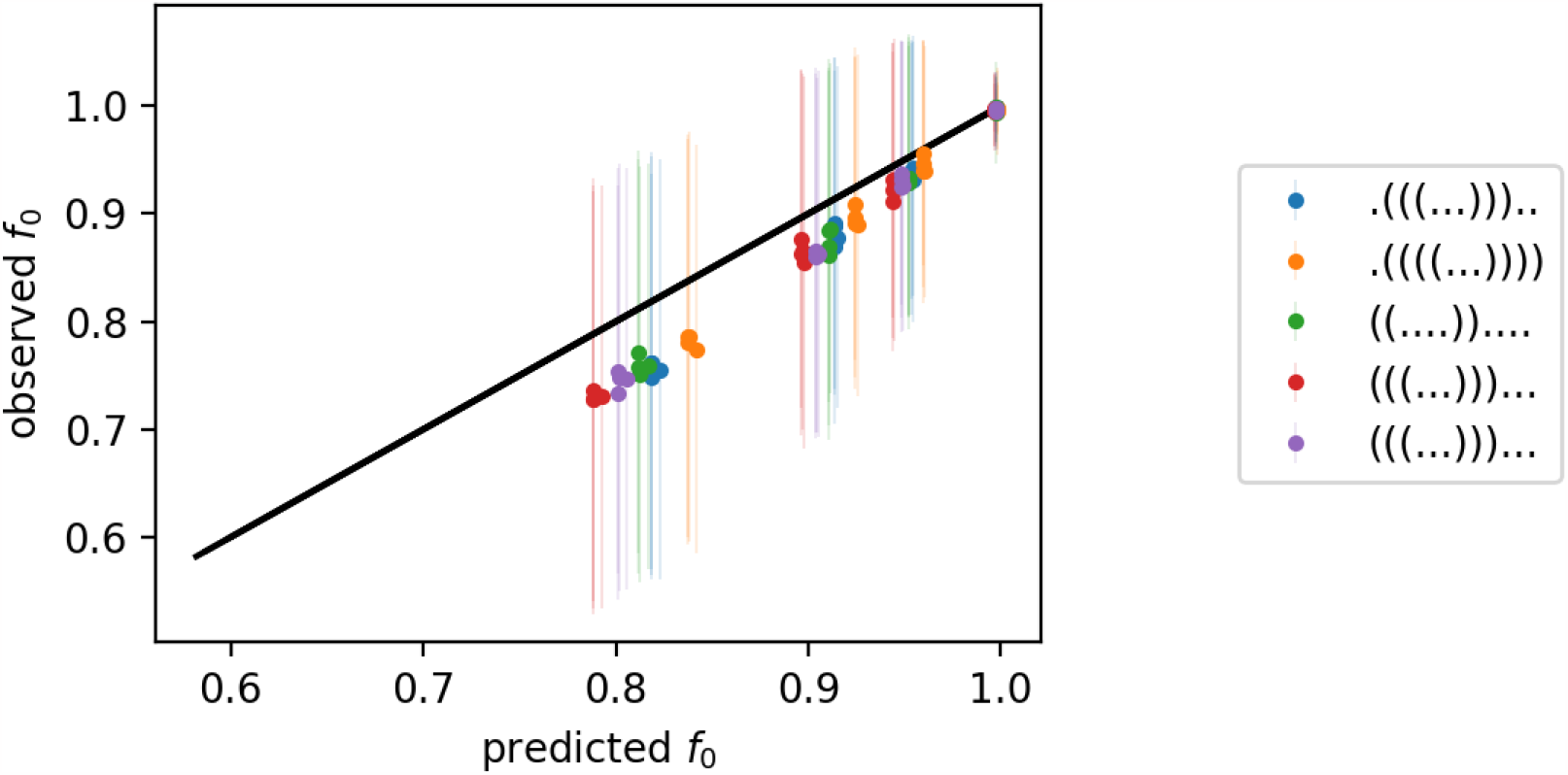
Comparing analytic estimates of the amount of polymorphism in a population to the simulation data: the figure shows both analytic estimates and simulation data for the fraction of individuals f_0_ that share the prevalent genotype. The simulations are based on the same NCs as Fig S2 above, and the corresponding phenotype is given in the legend. We use a range of population parameters, both monomorphic populations (LuN ≪ 1) and populations with more genetic diversity (LuN ≈ 0.5): population sizes N = 10, N = 100, N = 1000, N = 2000 and mutation rates u = 0.0005/N, u = 0.01/N, u = 0.02/N and u = 0.05/N. The simulations are run for 10^6^ generation, and f_0_ is measured every 10^3^ generations and averaged over; the error bars indicate the standard deviation in the measurements.

Fig S20 shows how this approximation compares to the measured value in simulations for a range of population parameters *u* and *N* . We find that the approximate values correlate with the simulation results and are of the right order of magnitude. This is “good enough” for our purposes, where we want to use a very simple treatment of the population in order to simplify the remaining populations. However, it is also clear that even in this example, the approximation is not perfect, and it is expected to break down for highly polymorphic populations, which we are not considering in this paper. Furthermore, the population diversity varies throughout the simulations, indicating that any single number cannot capture the full complexity. Therefore, it should not be applied to new situations without further tests.

### 5 Additional calculations

#### 5.1 Two exponential processes: which occurs first

In the main text, we needed the probability that an event of type 2 occurs before the first event of type 1. Let us assume that the two events 1 and 2 occur at constant rates, one at rate 1*/τ*_1_ and the other at rate 1*/τ*_2_. Then the probability that the next event of type 2 occurs before the next event of type 1, i.e. *t*_2_ *≤ t*_1_, is given by an integral over the corresponding exponential process (if we neglect the fact that time is measured in discrete generations):

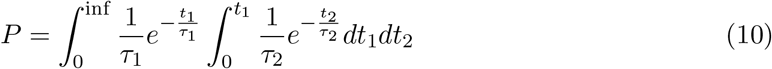

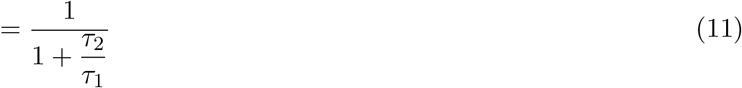

#### 5.2 Generalisation to more than two exponential processes: which occurs first

Let us generalise the calculation in the previous section to more than three types of events (for example the fixation of three different phenotypes): if we assume that all three events can be described as three independent exponential processes, then we need to calculate the probability that the next event of type 1 occurs before the next event of types 2 and 3, i.e. *t*_1_ *≤ min*(*t*_2_, *t*_3_). We can simplify this calculation by combining types 2 and 3, which occur at rates 1*/τ*_2_ and 1*/τ*_3_ respectively, into a single exponential process with rate (1*/τ*_2_ + 1*/τ*_3_) and time scale (1*/τ*_2_ + 1*/τ*_3_)^*−*1^. Then we can use Eq 11 to write down the probability that an event of type 1 occurs before the first event of the combined 2 & 3 process as:

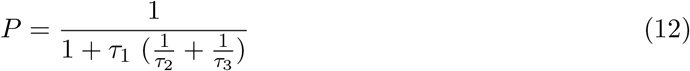

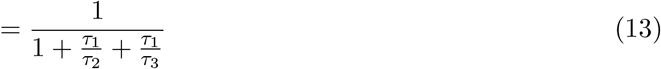

This expression can easily be generalised to *n* outcomes (for example if instead of having a two-peaked landscape, we have an n-peaked one). In this case, the rate of all events except type one is 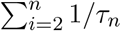, which means that the probability that an even of type 1 is observed first is given by:

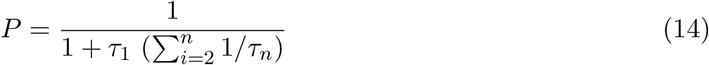

Note, however, that all this only holds while events can be approximated as independent exponential processes. This is not the case, for example, if *τ*_*n*_ stand for the appearance times of mutant *n* and several types of mutants are accessible from the same part of the NC.

#### 5.3 Two exponential processes: when do we expect the first event

Let us again assume that two types of events occur at constant rates, one at rate 1*/τ*_1_ and the other at rate 1*/τ*_2_. Then the total rate is given by

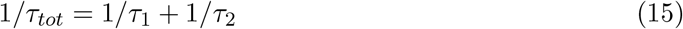

Therefore, the expected time until an event of *any* type is observed is given by:

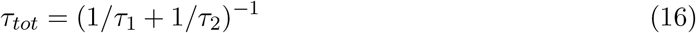

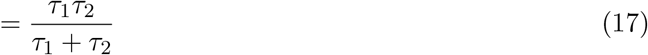

## Footnotes

1 Note that in our simple calculations we don’t distinguish between census size *N* and the effective population size *N*_*e*_, but these can be quite different [1]

2 We also confine the population to a single NC, not the entire neutral set of the initial phenotype. This restriction ensures comparability between our hierarchy of GP map models by preventing rare cases where the population moves to a different NC from the neutral set of *p*_*g*_ after a combination of a specific double mutation and genetic drift.

3 Strictly speaking a geometric distribution since time is measured in discrete generations.

